# Functional expression and sex dimorphism of the T-type Cav3.2 Calcium Channel in human DRG Neurons

**DOI:** 10.1101/2024.12.27.630548

**Authors:** Jean Chemin, Vanessa Soubeyre, Stephanie Shiers, Amaury François, Gaëtan Poulen, Nicolas Lonjon, Florence Vachiery-Lahaye, Luc Bauchet, Pierre François Mery, Theodore J Price, Emmanuel Bourinet

## Abstract

T-type/Cav3 calcium channels are key in neuronal excitability and pain processing with Cav3.2 being the prominent isoform in primary sensory neurons of the dorsal root ganglion (DRG). Its pharmacological inhibition or gene silencing induces analgesia in several preclinical models of inflammatory and neuropathic pain. However, the presence of Cav3.2, encoded by the *CACNA1H* gene, in human DRG neurons remains unresolved. Using RNA in-situ hybridization and electrophysiological recordings, we show that human DRGs express Cav3.2 in a subset of neurons positive for the neurotrophic factor receptor TrkB (*NTRK2* gene). The Cav3.2 current exhibits typical biophysical and pharmacological properties, including inhibition by a low concentration of nickel and by Z944, a specific T-type calcium channel blocker in advanced clinical development. Conversely, ABT-639, a T-type calcium channel inhibitor that failed in Phase 2 trials for pain relief, does not inhibit Cav3.2 currents in human DRG neurons. Importantly, Cav3.2 currents are prominent in neurons from female organ donors, supporting the presence of sex differences in pain mechanisms in humans. These findings underscore the potential of continued exploration of Cav3.2 as a therapeutic target for pain treatment and highlight a specific subset of human neurons that likely rely on this channel to modulate their excitability.

## Introduction

Chronic pain is a debilitating pathology with a very high prevalence, affecting hundreds of millions of people worldwide (1). Current treatments for chronic pain often have limited efficacy or life-threatening side effects, as exemplified by the opioid crisis in the USA (2). The need for new medications for chronic pain is therefore a critical medical, social and economic challenge. However, despite decades of extensive studies in animal models, few new therapeutic molecules have emerged. Therefore, understanding the molecular neurobiology of pain in humans is a prerequisite for the therapeutic translation of results obtained from animal studies (3, 4).

Sensory neurons in the DRG and trigeminal ganglion (TG) are the primary input for nociceptive signals and thus represent an attractive therapeutic target for painful conditions. Sensory neurons express a large panel of ion channels that contribute to their excitability and consequently to somatosensation, including pain (4, 5). Voltage-gated calcium channels (VGCCs) play an important role in the physiology of sensory neurons, contributing to both cellular excitability and neurotransmitter release (6). VGCCs include the high-voltage-activated (HVA) calcium channels (Cav1 and Cav2 families) and the low-voltage-activated (LVA) calcium channels (T-type / Cav3 family) (7). The LVA / T-type calcium channels are activated at voltages below the action potential threshold and thus contribute crucially to cellular excitability (8, 9). T-type calcium channels comprise three isoforms: Cav3.1, Cav3.2, and Cav3.3, among which Cav3.2 is expressed in rodent sensory neurons, including nociceptors and mechanoreceptors (9–14). Preclinical studies in rodents over the last decade have demonstrated the important role of Cav3.2 in both acute pain and several models of chronic pain of different etiologies (15–20). However, whether Cav3.2 is functionally expressed in human sensory neurons remains an open question (21). Furthermore, the role of Cav3.2 in human pain perception has been questioned since ABT-639, a peripherally restricted T-type calcium channel blocker (22), failed to alleviate pain in patients with diabetic peripheral neuropathic pain (23–25). In order to translate the results obtained in rodents to humans, we investigated here the expression, function, and the pharmacology of the T-type calcium channels in sensory DRG neurons obtained from organ donors.

## Results

The presence of T-type calcium channels was first analyzed by in-situ hybridization using RNAscope quantification with chromogenic and fluorescent dyes. We first investigated the presence of CACNA1G (Cav3.1), CACNA1H (Cav3.2), and CACNA1I (Cav3.3) mRNAs, in lumbar and thoracic DRG sections from 7 organ donors (3 males and 4 females, aged 36 to 75 years (58 ± 5.2 years old, Table 1) using a chromogenic approach (Figure 1). In these experiments, DRG cells were identified by haematoxylin staining (Figure 1A), and the control quality of mRNAs labeling was confirmed using the manufacturer’s positive and negative probes (Supplementary Figure 1A and 1B). We also confirmed the identity of visually identified human DRG neurons using a probe targeting synaptotagmin 1 mRNA (Syt 1 gene, Supplementary Figure 1C-D), which has been reported as a pan-neuronal marker in human DRG single-cell transcriptomic databases (26, 27). We found that Cav3.2 mRNA, encoded by the *CACNA1H* gene, was robustly expressed with ∼60 individual RNA dots per cell in 16% of DRG neurons (Figure 1A-C), as well as in blood vessels (Supplementary Figure 1E). Cav3.2 mRNA expression was similar in DRGs from both male and female organ donors (Supplementary Figure 1F), indicating no sex bias at the RNA level. Cell diameter analysis revealed that Cav3.2 mRNA was expressed in DRG neurons of medium diameter, with a peak in neurons of 40-50 µM diameter (Supplementary Figure 1G-H). In contrast, Cav3.1 mRNA was weakly expressed, with one large spot present exclusively in the nucleus (Figure 1D-F), whereas Cav3.3 mRNA showed limited expression in human DRGs, with less than 5 RNA puncta detected in a handful of neurons within each sample (Figure 1G-I). We next examined whether Cav3.2 mRNA was coexpressed with Nav1.8 mRNA, a marker of nociceptors in rodents and humans (26, 27). Nav1.8 mRNA (SCN10A gene) was highly expressed in a large population of human DRG neurons (∼60 %, Figure 1) with small cell diameter (∼38 µM, Supplementary Figure 1G-H). Strikingly, most Cav3.2 mRNA-positive neurons (92 %) did not express Nav1.8 mRNA (Figure 1B-C), whereas the opposite pattern was found for Cav3.1 mRNA, which was predominantly expressed in Nav1.8-positive neurons (Figure 1E-F). Similar results were found in another set of experiments performed independently using the RNAscope fluorescence approach in a different cohort of donors with a different cause of death (2 males and 2 females, aged 19 to 75 years (38.3 ± 12.5 years, Table 1) using RNAscope fluorescent dyes for Cav3.1, Cav3.2, and Cav3.3 mRNA (Figure 2). In these experiments, Cav3.2 mRNA was again robustly expressed in a subset of medium diameter neurons (Figure 2C and 2D), as well as in blood vessels (Figure 2K orange arrows), whereas Cav3.1 mRNA was weakly expressed and similarly restricted to the nucleus of human DRG neurons (Figure 2A and 2B). Results for Cav3.3 mRNA were also independently confirmed showing limited expression in very few neurons within each sample (Figure 2E and 2F). On average, these experiments showed that Cav3.2 was expressed in 17.1 %, Cav3.1 in 6.3 % and Cav3.3 in 1.6 % of the total neuronal population (Figure 2G). We also found that TRPV1 mRNA, another marker of nociceptive neurons in rodents and humans (26, 27), was highly expressed in 80 % of the neurons (Figure 2G). As previously obtained with Nav1.8, Cav3.2 RNA was enriched in TRPV1-negative neurons (Figure 2C and 2I) and weakly expressed in TRPV1-positive neurons (Figure 2I), whereas an opposite pattern was observed for Cav3.1 RNA (Figure 2A and 2H).

**Figure 1.**
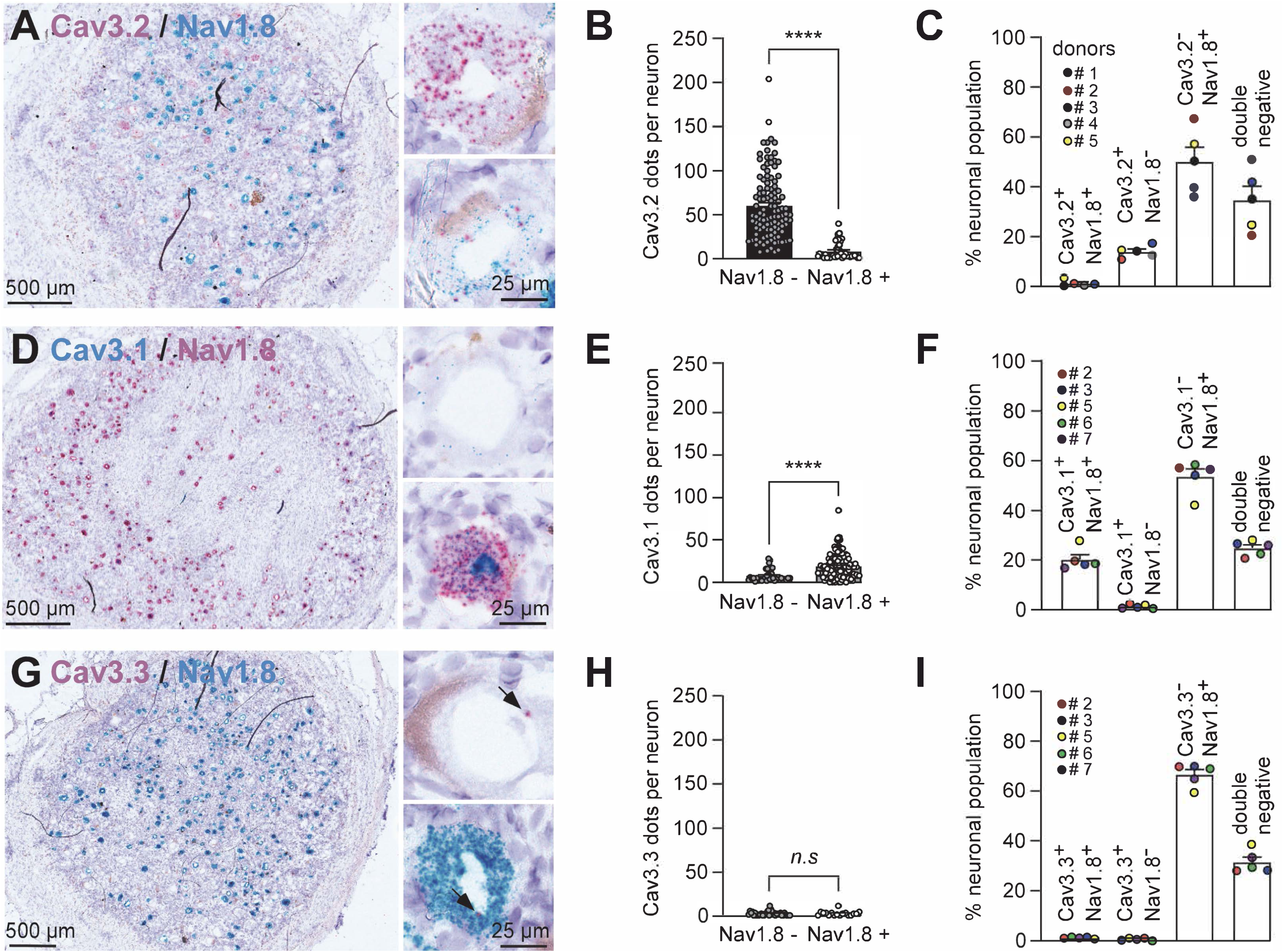
Distribution of Cav3 and Nav1.8 mRNA in human dorsal root ganglia. (**A**) Representative images of human DRGs (hDRGs) labeled with RNAscope chromogenic *in situ* hybridization for Cav3.2 (pink) and Nav1.8 (gene *SCN10A*, blue) with haematoxylin (purple). Magnification showing a Cav3.2^+^/Nav1.8^-^ neuron (upper inset) and a Cav3.2^+^/Nav1.8^+^ neuron (lower inset). **(B)** Comparison of the number of Cav3.2 mRNA dots in Nav1.8-negative and Nav1.8-positive neurons (non-parametric two-tailed Mann-Whitney test, *p*<0.0001). **(C)** Subpopulation distribution of Cav3.2 in combination with Nav1.8 in the hDRG. **(D)** Representative image of hDRGs labeled with RNAscope chromogenic *in situ* hybridization for Cav3.1 (blue) and Nav1.8 (pink) with haematoxylin (purple). Magnification a Cav3.1^+^/Nav1.8^-^ neuron (upper inset) and a Cav3.1^+^/Nav1.8^+^ neuron (lower inset). **(E)** Comparison of the number of Cav3.1 mRNA dots in Nav1.8-negative and Nav1.8-positive neurons (non-parametric two-tailed Mann-Whitney test, *p*<0.0001). **(F)** Subpopulation distribution of Cav3.1 in combination with Nav1.8 in the hDRG. **(G)** Representative image of hDRGs labeled with RNAscope chromogenic *in situ* hybridization for Cav3.3 (pink) and Nav1.8 (gene SCN10A, blue) with haematoxylin (purple). Magnification showing a Cav3.3^+^/Nav1.8^-^ neuron (upper inset) and a Cav3.3^+^/Nav1.8^+^ neuron (lower inset). **(H)** Comparison of the number of Cav3.3 mRNA dots in Nav1.8-negative and Nav1.8-positive neurons (non-parametric two-tailed Mann-Whitney test, *p*=0.95). **(I)** Subpopulation distribution of Cav3.3 in combination with Nav1.8 in the hDRG.

**Figure 2.**
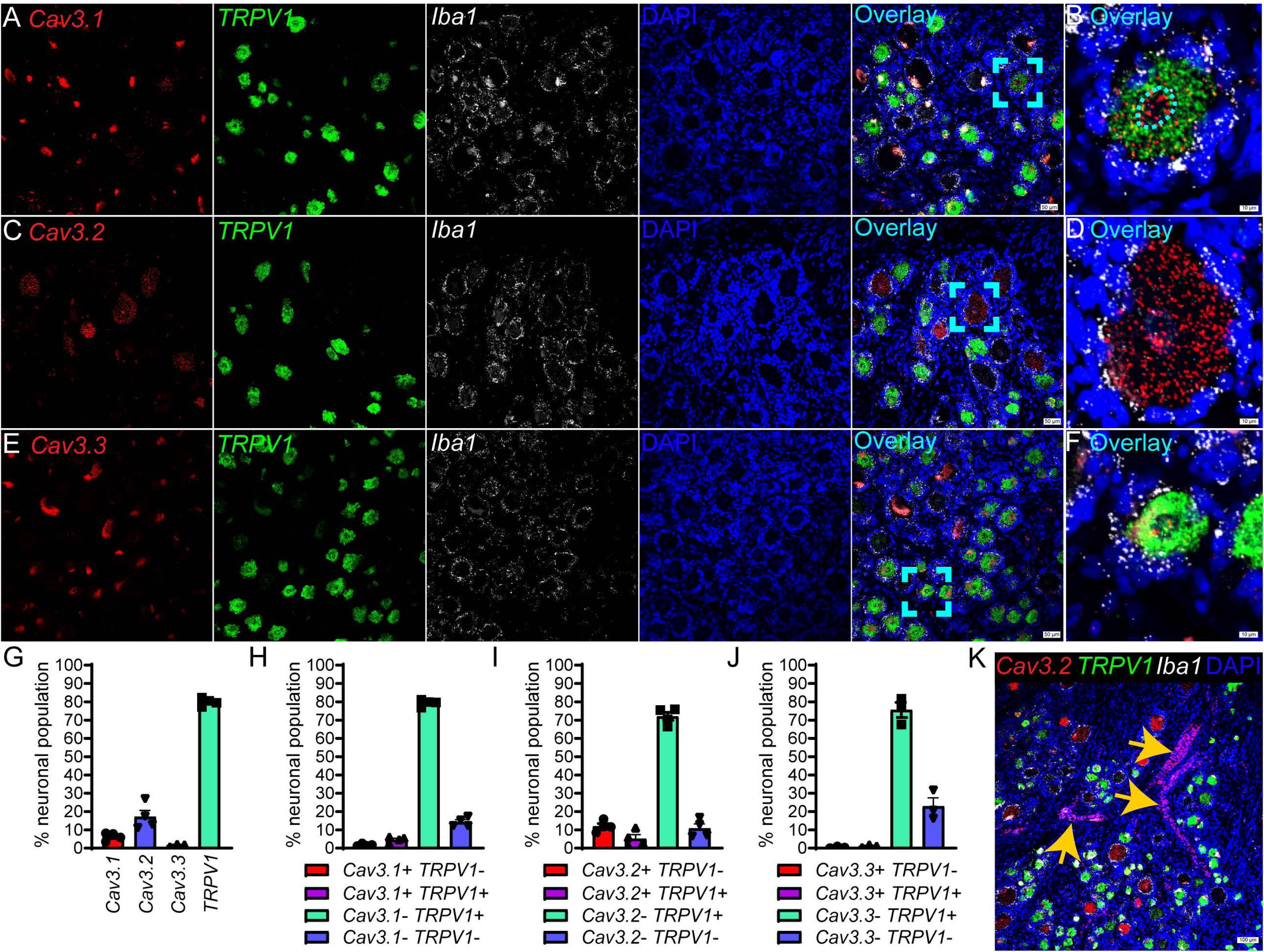
Distribution of Cav3 and TRPV1 mRNA in human dorsal root ganglia. **(A)** Representative images of hDRGs labeled with RNAscope fluorescent *in situ* hybridization for Cav3.2 (gene CACNA1H, pink), TRPV1 (green), Iba1 (gene AIF1, white), and DAPI (blue). The cyan outline in the overlay image marks a **(B)** Cav3.1+ sensory neuron. Cav3.1 mRNA was lowly expressed in a small subset of sensory neurons and showed enrichment within the nucleus (cyan outline). **(C)** Representative images of human lumbar DRGs labeled with RNAscope fluorescent *in situ* hybridization for Cav3.2 *(gene CACNA1H,* red), TRPV1 (green), Iba1 (gene AIF1, white), and DAPI (blue). The cyan outline in the overlay image marks a **(D)** Cav3.2+ sensory neuron. **(E)** Representative images of human lumbar DRGs labeled with RNAscope fluorescent *in situ* hybridization for Cav3.3 *(gene CACNA1I,* red), TRPV1 (green), Iba1 (gene AIF1, white), and DAPI (blue). The cyan outline in the overlay image marks a **(F)** Cav3.3+ sensory neuron. **(G)** Distribution of Cav3.1, Cav3.2, Cav3.3 and TRPV1 in the total neuronal population. **(H)** Subpopulation distribution of TRPV1 in combination with Cav3.1, **(I)** Cav3.2, and **(J)** Cav3.3 in the hDRG. **(K)** Cav3.2 was also found in blood vessels (orange arrows). Scale bars = 50 μm in panels A, C, E. Scale bars = 10 μm in panels B, D, F. Scale bar = 100 μm in panel J. Large globular structures found in neurons are lipofuscin and were not considered as RNA signal.

**Table 1.** Characteristics of donors and experiments.

| Donor | Age | Sex | Cause of death | Experiments |
| --- | --- | --- | --- | --- |
| 1 | 53 | M | CVA/brain death | RNAscope chromogenic (Fig. 1) |
| 2 | 50 | F | aneurysm rupture//brain death | RNAscope chromogenic (Fig. 1 & 3) |
| 3 | 53 | M | CVA/brain death | RNAscope chromogenic (Fig. 1) |
| 4 | 68 | F | CVA/brain death | RNAscope chromogenic (Fig. 1)<br>Electrophysiology |
| 5 | 75 | F | CVA hemorrhagic /brain death | RNAscope chromogenic (Fig. 1 & 3) |
| 6 | 71 | M | CVA/brain death | RNAscope chromogenic (Fig. 1 & 3) /<br>Electrophysiology |
| 7 | 36 | F | CVA/brain death | RNAscope chromogenic (Fig. 1 & 3) |
| 8 | 29 | F | Anoxia/Drug Overdose | RNAscope fluorescent (Fig. 2) |
| 9 | 19 | M | Anoxia/Drug Overdose | RNAscope fluorescent (Fig. 2) |
| 10 | 30 | M | Anoxia/Asthma Attack | RNAscope fluorescent (Fig. 2) |
| 11 | 75 | F | CVA/Stroke | RNAscope fluorescent (Fig. 2) |
| 12 | 77 | F | brain death | RNAscope chromogenic (Fig. 3)<br>Electrophysiology |
| 13 | 76 | F | CVA hemorrhagic /brain death | RNAscope chromogenic (Fig. 3)<br>Electrophysiology |
| 14 | 68 | M | CVA hemorrhagic /brain death | RNAscope chromogenic (Fig. 3) |
| 15 | 76 | M | brain death | Electrophysiology |
| 16 | 56 | F | CVA/brain death | Electrophysiology |
| 17 | 71 | M | CVA hemorrhagic /brain death | Electrophysiology |
| 18 | 33 | F | Arteriovenous malformation rupture/ brain death | Electrophysiology |
| 19 | 52 | M | aneurysm rupture//brain death | Electrophysiology |
| 20 | 46 | F | CVA hemorrhagic /brain death | Electrophysiology |

The presence of Cav3.2 RNA was further investigated in combination with those of TrkB (*NTRK2* gene) and P2Y1 (*P2RY1* gene), two low-threshold mechanoreceptor-specific transcripts, as well as with TRPM8 RNA (the prototypical cold sensor marker) in thoracic and lumbar DRG sections from 7 organ donors (2 males and 5 females, aged 36 to 77 years (64.7 ± 5.9 years, Figure 3). Cav3.2 was widely expressed in 60 % of TrkB-positive neurons (Figure 3A-B), which represented 30 % of the total neuronal population (Figure 3B) and had a mean diameter of ∼58 µM (Figure 3C and Supplementary Figure 1H). Thus, approximately 86% of Cav3.2-positive neurons expressed TrkB (Figure 3B). In addition, both Cav3.2 and TrkB were also co-expressed in blood vessels (Supplementary Figure 1E). The Cav3.2 mRNA was also expressed in 45 % of P2Y1-positive neurons (Figure 3D-E), which represented 32 % of the total neuronal population (Figure 3E) and had a mean diameter of 44 µM (Figure 3F and Supplementary Figure 1H). In addition, Cav3.2 was also expressed in a subset of TRPM8-positive neurons (∼17 %, Figure 3G-H). The TRPM8 positive population in these human DRG samples represented 24 % of the total neuronal population (Figure 3H) and had a mean diameter of 40 µM (Figure 3I and Supplementary Figure 1H). In these neurons, both Cav3.2 and TRPM8 showed much lower expression as compared to TRPM8^+^/Cav3.2^-^ and Cav3.2^+^/TRPM8^-^ expressing neurons (Supplementary Figure 1I). Overall, RNAscope experiments indicated the presence of the Cav3.2 RNA in approximately 18 % of neurons, ranging in diameter from 25 to 85 µM, with a peak expression in neurons of 40-50 µM diameter.

**Figure 3.**
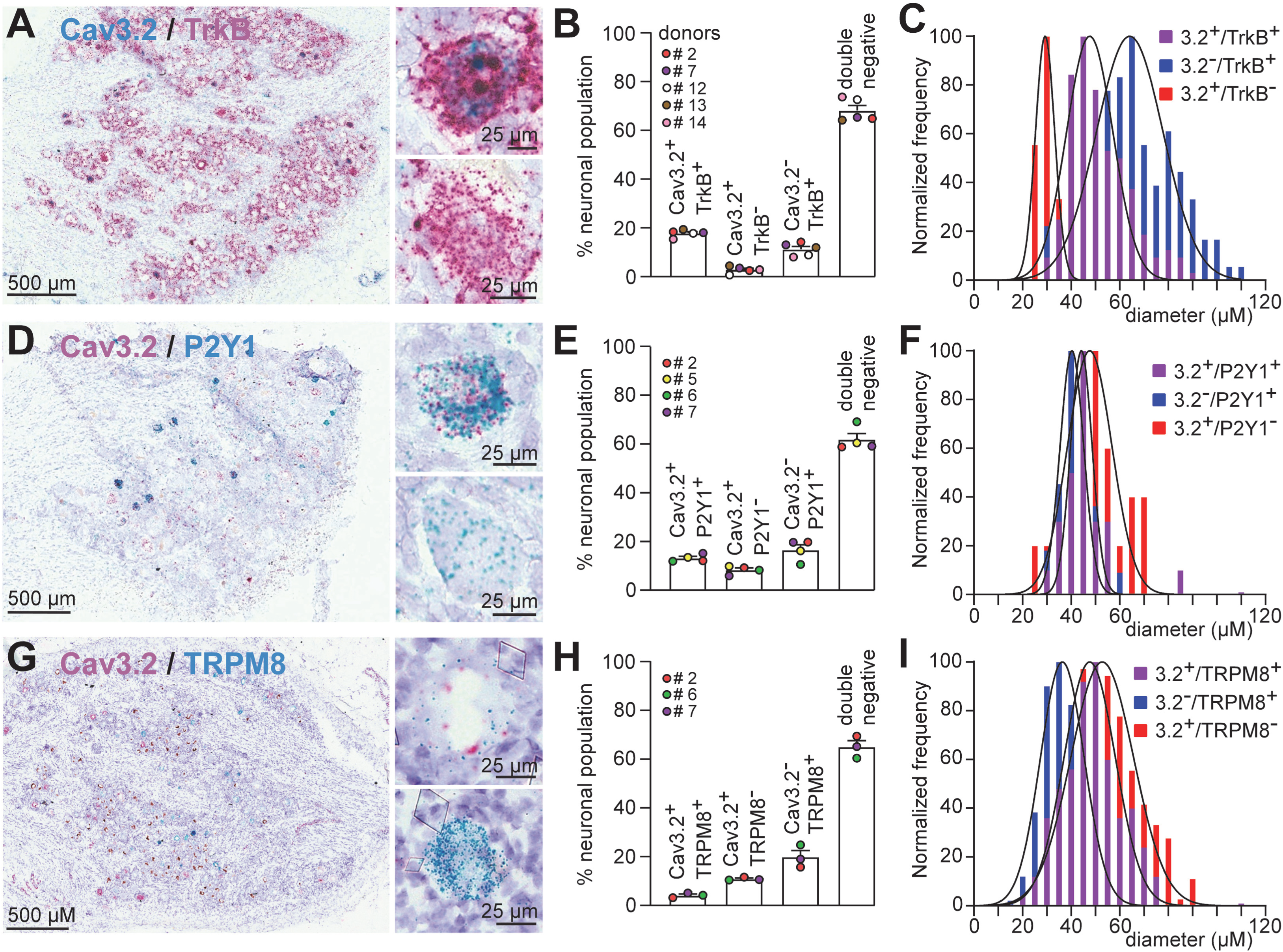
Distribution of Cav3.2 with different markers of human DRG neurons. **(A)** Representative image of hDRGs labeled with RNAscope chromogenic *in situ* hybridization for Cav3.2 (blue) and TrkB (gene *NTRK2*, pink) with haematoxylin (purple). Magnification of the DRG slice showing a Cav3.2^+^/TrkB^+^ neuron (upper inset) and a Cav3.2^-^/TrkB^-^ neuron (lower inset). **(B)** Subpopulation distribution of Cav3.2 in combination with TrkB in the hDRG. **(C)** Normalized frequency (%) of hDRG neurons expressing Cav3.2 and/or TrkB relative to cell diameter. **(D)** Representative image of hDRGs labeled with RNAscope chromogenic *in situ* hybridization for Cav3.2 (pink) and P2Y1 (gene *P2RY1*, blue) with haematoxylin (purple). Magnification of the DRG slice showing a Cav3.2^+^/P2Y1^+^ neuron (upper inset) and a Cav3.2^-^/P2Y1^-^ neuron (lower inset). **(E)** Subpopulation distribution of Cav3.2 in combination with P2Y1 in the hDRG. **(F)** Normalized frequency of hDRG neurons expressing Cav3.2 and/or P2Y1 relative to cell diameter. **(G)** Representative image of human DRGs labeled with RNAscope chromogenic *in situ* hybridization for Cav3.2 (pink) and TRPM8 (blue) with haematoxylin (purple). Magnification of the DRG slice showing a Cav3.2^+^/TRPM8^+^ neuron (upper inset) and a Cav3.2^-^/TRPM8^-^ neuron (lower inset). **(H)** Subpopulation distribution of Cav3.2 in combination with TRPM8 in the hDRG. **(I)** Normalized frequency of hDRG neurons expressing Cav3.2 and/or TRPM8 relative to cell diameter.

Electrophysiological investigation of the presence of the T-type calcium current in human sensory neurons was performed in cultures of thoracic and lumbar DRG neurons from 10 organ donors (6 females and 4 males, aged 33 to 77 years (62.6 ± 4.7 years old, Table 1). A double-pulse protocol (P/8 leak subtraction protocol), consisting of a small depolarization at – 40 mV (allowing the activation of the LVA / T-type current) and a stronger depolarization at 0 mV (allowing the activation of the HVA current), was performed from a holding potential of – 90 mV (Figure 4). Using this protocol (Figures 4A and 4C), we recorded robust HVA calcium currents in neurons from all donors (N=10 donors, n=101 neurons) and a LVA / T-type calcium current, of smaller amplitude, in a subset (∼20 %) of DRG neurons (Figure 4A, N=7 donors, n=22 neurons). In these neurons, the LVA / T-type calcium current exhibited distinct kinetics than the HVA current, including slower activation, faster inactivation and slower deactivation (Figures 4A and 4B, n=22). After 100ms, the amplitude of the T-type current decreased by approximately 70 %, while the amplitude of the HVA current decreased by only approximately 20 % (Figure 4A). On average, the amplitude of the LVA current was ∼1 nA (Figure 4D) and the current density was ∼10 pA/pF (Figure 4E), while those of the HVA current were ∼5 nA and ∼50 pA/pF, respectively. In the remaining neurons (∼80 %), only HVA current was observed (Figure 4C, n=79). Strikingly, HVA current amplitude and density were lower in these neurons (Figure 4D and 4E), and a correlation between LVA current and HVA current was observed when all neurons were analyzed (Supplementary Figure 2B). In contrast, no difference in cell diameter (∼45 µM) or in cell membrane capacitance (∼100 pF) was observed with respect to T-type current expression (Figure 4C, inset), whereas cell diameter was positively correlated with membrane capacitance (Supplementary Figure 2A). T-type current expression was preferentially observed in neurons categorized by a diameter range of 40-50 µM (Supplementary Figure 2C) and a membrane capacitance range around 80 pF (Supplementary Figure 2D). Interestingly, in contrast to the HVA current, the amplitude of the T-type current was not positively correlated with membrane capacitance or cell diameter (Supplementary Figure 2E-H).

**Figure 4.**
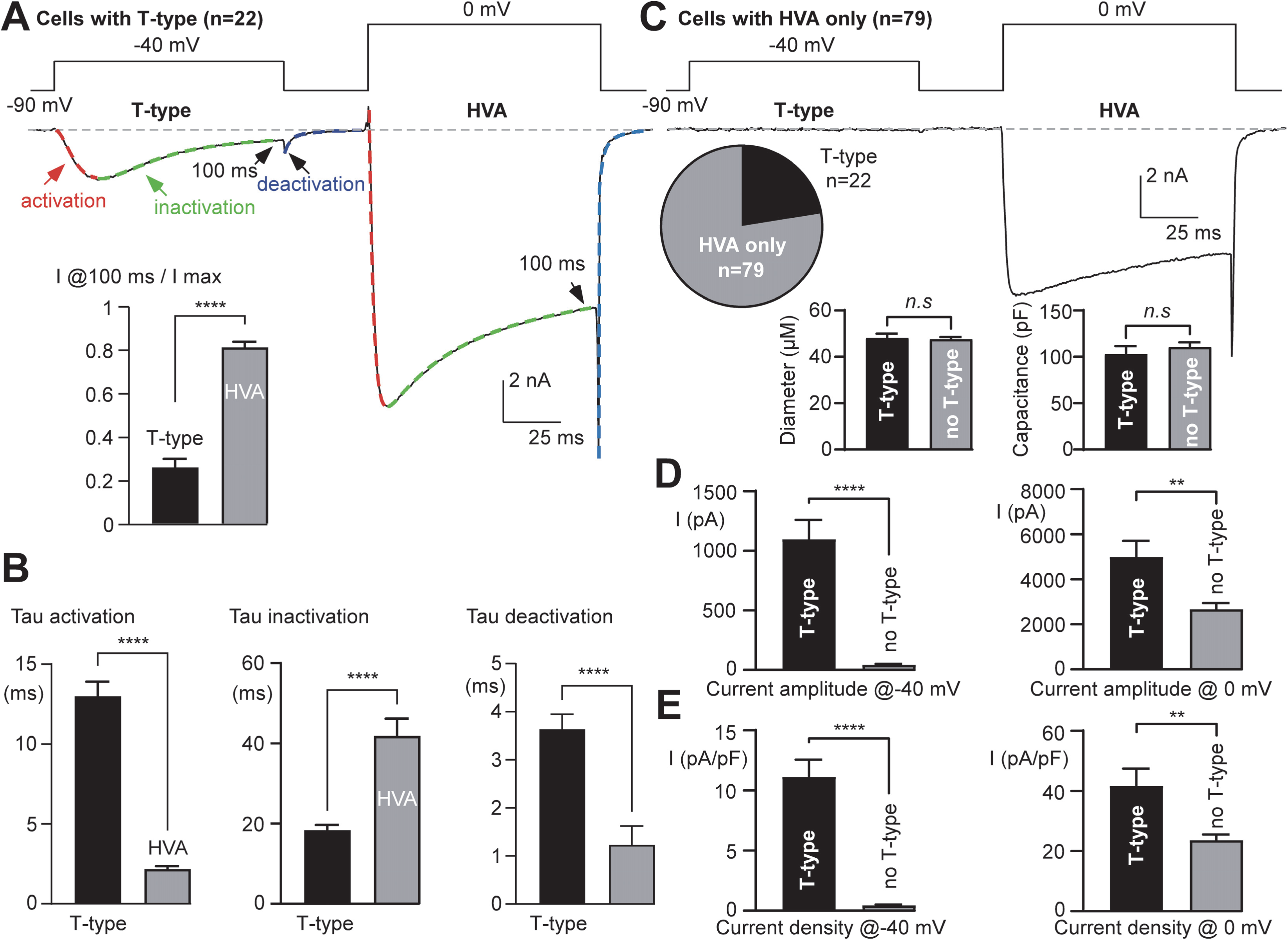
Expression of a LVA / T-type calcium current in human DRG neurons. **(A)** Exemplar neuron expressing two types of calcium current elicited by a double-pulse protocol (LVA at –40 mV and HVA at 0 mV). Amplitude of the current remaining after 100 ms normalized to the maximal current (inset, non-parametric two-tailed Mann-Whitney test, *p*<0.0001, n=22). **(B)** Current kinetics of the LVA / T-type calcium current (elicited at –40 mV) compared with those of the HVA calcium current (elicited at 0 mV) (non-parametric two-tailed Mann-Whitney test, n=22, τ activation *p*<0.0001, τ inactivation *p*<0.0001, and τ deactivation *p=*0.0001). **(C)** Exemplar neuron expressing only HVA calcium current. Pie chart showing the proportion of neurons expressing T-type calcium current (∼20 %, n=101). Comparison of the diameter and membrane capacitance of neurons expressing and not expressing T-type calcium current (inset, n=101, non-parametric two-tailed Mann-Whitney test, diameter *p*=0.937, membrane capacitance *p*=0.499). **(D)** Current amplitude measured at –40 mV (left, *p*<0.0001) and at 0 mV (right, *p*=0.0011) in neurons expressing (n=22) and not expressing (n=79) T-type calcium current (non-parametric two-tailed Mann-Whitney test). **(E)** Current density measured at –40 mV (left, *p*<0.0001) and at 0 mV (right, *p*=0.0002) in neurons expressing (n=22) or not expressing (n=79) T-type calcium current (non-parametric two-tailed Mann-Whitney test).

The steady-state activation properties of the T-type calcium current were then examined using increasing step depolarizations (Figure 5). When applied from a holding potential (HP) of –90 mV, depolarizations between –60 mV and –30 mV induced a progressive activation of the T-type calcium current (Figure 5A, black traces), which peaked around –30 mV (Figure 5A and 5B). In contrast, stronger depolarizations were required to activate the HVA calcium current, which peaked around –10 mV (Figure 5A and 5B, gray traces). In cells expressing the T-type calcium current, the current-voltage (IV) curve was fitted with two components to account for the presence of LVA and HVA calcium currents (Figure 5B). This analysis revealed that the T-type calcium current developed with a half-activation potential (V0.5) of –45.96 ± 1.41 mV and a Boltzmann slope of –4.39 ± 0.33 (n=16), whereas the V0.5 and slope values for the HVA calcium current were –16.46 ± 0.33 mV and –2.35 ± 0.36 (n=16), respectively. Because the T-type calcium current was rapidly inactivated, the IV curve obtained when currents were measured at 100 ms indicated the presence of only the HVA calcium current (Figures 5A and 5B). As observed previously, the maximal HVA calcium current density was lower in cells not expressing the T-type calcium current (30.23 ± 2.79 pA/pF (n=57) vs 55.42 ± 9.33 pA/pF (n=16) in cells expressing the T-type calcium current, p=0.011, non-parametric two-tailed Mann-Whitney test, Figure 5B). In addition, the IV curve obtained from HP –60 mV (Figure 5C and 5D) indicated the presence of the HVA calcium current only, and at this HP no current was recorded at –30 mV in cells expressing T-type calcium current (Figure 5C). The dependence of the T-type calcium current on the HP was examined in more detail at HPs ranging from –110 mV to – 40 mV (Figure 5E). The T-type calcium current showed 14.5 ± 3.3 % inactivation (n=11) at HP –90 mV, 37.5 ± 5.2 % inactivation (n=11) at HP –80 mV, 62.9 ± 4.9 % inactivation (n=11) at HP –70 mV and 92.8 ± 1.9 % inactivation (n=11) at HP –60 mV (Figure 5E and 5F). The steady-state inactivation curve was fitted with a Boltzmann equation (Figure 5F), indicating a half inactivation potential (V0.5) of –75.97 ± 1.74 mV and a slope of 7.05 ± 0.59 (n=11). Interestingly, the overlap of the inactivation and activation curves indicated the existence of a window current at physiological resting potential around –70 to –50 mV (Figure 5F).

**Figure 5.**
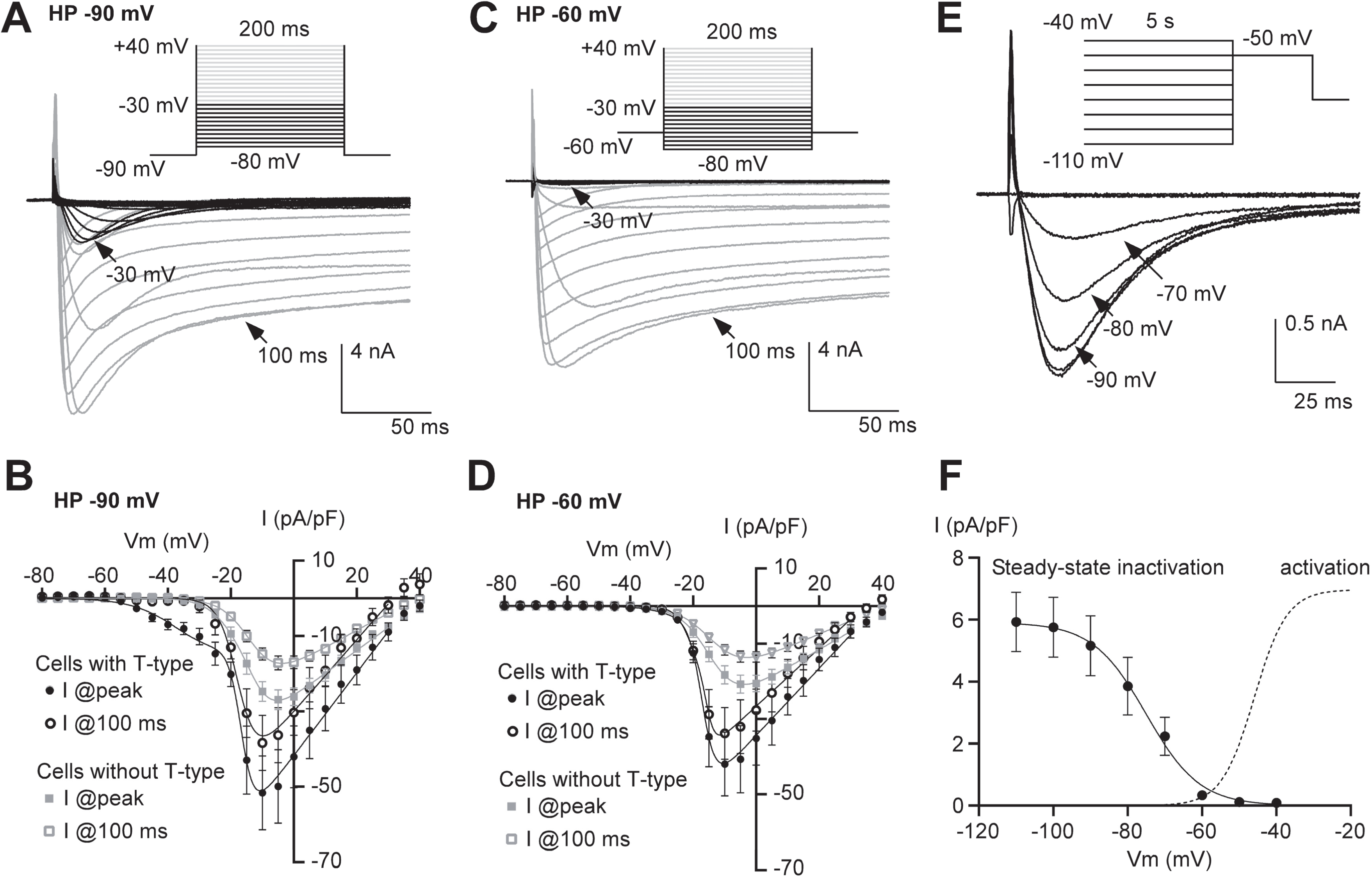
Activation and inactivation properties of T-type calcium current in human DRG neurons. (**A**) Exemplar calcium currents elicited by a series of step depolarizations ranging from −80 to +40 mV (0.2 Hz stimulation frequency) from a holding potential (HP) of −90 mV. Note the presence of the T-type calcium current elicited by depolarizations below –30 mV (black traces). (**B**) Current-voltage (I-V) curves of the calcium current obtained in cells expressing (black circles, n=16) or not expressing (gray squares, n=57) the T-type calcium current. Currents were measured at the peak (filled symbols) or at 100 ms (empty symbols) from an HP of –90 mV. (**C**) Exemplar calcium currents elicited by a series of step depolarizations ranging from −80 to +40 mV (0.2 Hz stimulation frequency) from a holding potential of −60 mV (same neuron as in A). Note the absence of the LVA / T-type calcium current elicited by depolarizations below –30 mV. **(D)** Current-voltage (I-V) curves of the calcium current obtained in cells expressing (black circles, n=11) or not expressing (gray squares, n=39) the T-type calcium current. Currents were measured at the peak (filled symbols) or at 100 ms (empty symbols) from an HP of –60 mV. **(E)** Steady-state inactivation of the T-type calcium current. The T-type calcium current was elicited at –50 mV from a series of HPs ranging from −110 to –40 mV (0.2 Hz stimulation frequency). **(F)** Steady-state inactivation curve of the T-type calcium current (n=11) and activation curve extracted from the I-V curve fitting shown in B (cells with T-type, peak in pA/pF). Note the presence of a window current resulting from the overlap of the inactivation and activation curves.

The pharmacological properties of the T-type calcium current were then investigated using cadmium (a general calcium channel blocker (28), nickel, at low concentration (which mainly inhibits Cav3.2 (29), and Z944 (a specific T-type calcium channel inhibitor (30) in an ongoing Phase 3 clinical trial for essential tremor (#NCT06087276; ClinicalTrials.gov). Application of 200 µM cadmium inhibited both T-type and HVA calcium currents recorded using the double-pulse protocol (Figure 6A). This inhibition occurred in the minute range and was reversible upon washout of the cadmium solution (Figure 6A inset). On average, 200 µM cadmium inhibited the T-type calcium current by 86.6 ± 2.2 % (n=7) and the HVA calcium current by 98.5 ± 0.6 % (n=22, Figure 6B). Similar results were observed during the I-V protocol, where cadmium inhibited both currents at all potentials (Figure 6C). In contrast, the application of 30 µM nickel mainly inhibited the T-type calcium current with a weak effect on the HVA calcium current (Figure 6D). The effects of nickel also occurred rapidly and were completely reversed after washout of the nickel solution (Figure 6D inset). Application of nickel resulted in inhibition of the T-type calcium current by 80.8 ± 2.6 % (n=12), while the HVA calcium current was inhibited by only 14.0 ± 1.8 % (n=25) (Figure 6E). During the I-V protocol, nickel perfusion inhibited the calcium current recorded at potentials between –50 to –25 mV, with a weak effect on the calcium current recorded at higher potentials (Figure 6F), indicating that the effect of nickel in human sensory neurons was selective for the T-type calcium current. The effect of the specific T-type calcium channel blocker, Z944, was then investigated (Figure 6G-I). Application of 1 µM Z944 inhibited the T-type calcium current by 75.3 ± 3.5 % (n=8), while the HVA calcium current was inhibited by only 10.2 ± 3.4 % (n=14, Figure 6H). The effect of Z944 developed within 1-2 minutes and was washed out within a few minutes during perfusion with the control solution (Figure 6G, inset). As observed with nickel during the I-V protocol, Z944 inhibited the calcium current recorded at low voltages with sparse effects on the calcium current recorded at higher voltages (Figure 6I). Overall, the pharmacological profile of T-type calcium current in human DRG neurons was similar to that observed in rodents (11, 16, 31–34) and indicated that Cav3.2 was the major isoform functionally expressed in these neurons. We also found that ABT-639, a previously described Cav3 inhibitor (22), did not inhibit the T-type calcium current in human DRG neurons (Figure 6J), and this result was confirmed using HEK-293 cells expressing recombinant hCav3.2 channels (Figure 6K). In addition to the typical nickel-sensitive T-type calcium current, we also found in a very small number of neurons (∼2 %, n=2 neurons out of 103) an atypical LVA current of low amplitude that was distinct from the T-type calcium current (Supplementary Figure 3). In these two neurons, the atypical LVA current activated more rapidly (Supplementary Figure 3D), inactivated more slowly (Supplementary Figure 3E), and peaked at more negative potentials than the T-type calcium current (at –45 mV, Supplementary Figure 3G-I). Importantly, this atypical LVA current was resistant to the application of nickel or Z944 (Supplementary Figure 3J and 3K). Therefore, these two cells were considered as neurons without T-type calcium current expression and were excluded from the previous and subsequent analyses.

**Figure 6.**
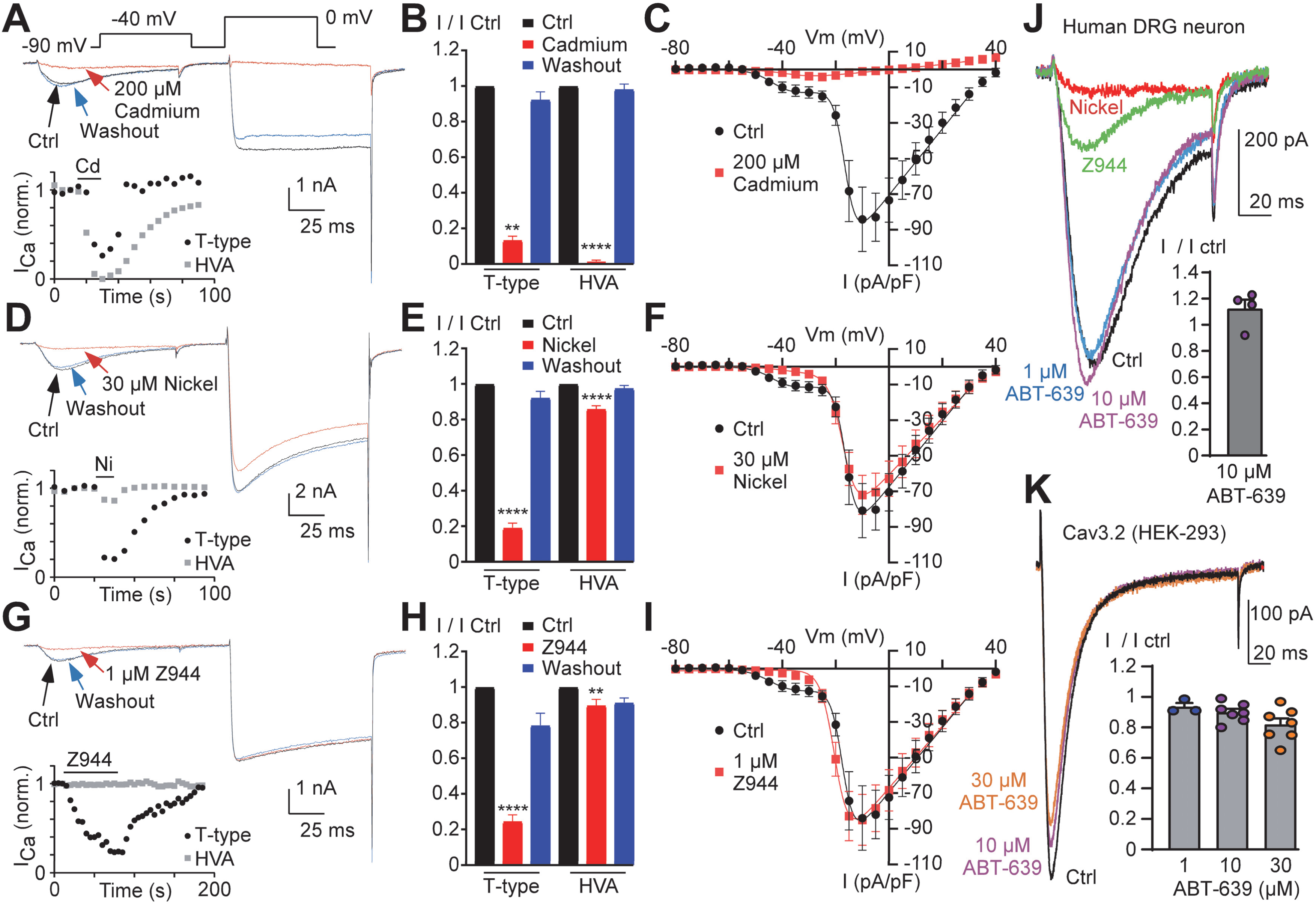
Pharmacological properties of T-type calcium current. (**A-C**) Inhibition of T-type and HVA calcium currents by cadmium. **(A)** Effect of a 200 µM cadmium perfusion and its washout on calcium currents elicited by the double-pulse protocol (0.2 Hz stimulation frequency). The time course of the current inhibition measured at –40 mV (T-type) and 0 mV (HVA) is presented as an inset. **(B)** Summary of the effect of 200 µM cadmium and its washout on the T-type (*p*=0.0015, n=7) and HVA (*p*<0.0001, n=22) calcium currents (non-parametric Friedman test with Dunn’s correction). **(C)** Current-voltage (I-V) curves in the presence and absence of cadmium (n=5). **(D-F)** Inhibition of T-type and HVA calcium currents by nickel. **(D)** Effect of a 30 µM nickel perfusion and its washout on calcium currents elicited by the double-pulse protocol (0.2 Hz stimulation frequency). The time course of the current inhibition measured at –40 mV (T-type) and 0 mV (HVA) is presented as an inset. **(E)** Summary of the effect of 30 µM nickel and its washout on T-type (n=12, *p*<0.0001) and HVA (*p*<0.0001, n=29) calcium currents (non-parametric Friedman test with Dunn’s correction). **(F)** Current-voltage (I-V) curves in the presence and absence of nickel (n=6). **(G-I)** Inhibition of T-type and HVA calcium currents by Z944. **(G)** Effect of a 1 µM Z944 perfusion and its washout on calcium currents elicited by the double-pulse protocol (0.2 Hz stimulation frequency). The time course of the current inhibition measured at –40 mV (T-type) and 0 mV (HVA) is presented as an inset. **(H)** Summary of the effect of 1 µM Z944 and its washout on T-type (*p*=0.0003, n=8) and HVA (*p*=0.0028, n=14) calcium currents (non-parametric Friedman test with Dunn’s correction). **(I)** Current-voltage (I-V) curves in the presence and absence of Z944 (n=5). **(J-K)** Lack of effect of ABT-639 at micromolar concentration on both T-type calcium current expressed in hDRG neurons (J, n=4) and human recombinant Cav3.2 current in HEK-293 cells (K, n=7). The currents were elicited at –40 mV from HP –90 mV (J) and at –30 mV from –80 mV (K).

The results obtained in individual organ donors were analyzed according to their age and sex (Figure 7). Strikingly, whereas the analysis of the occurrence and amplitude of the T-type calcium current showed a non-significant association with age (Figure 7A and Supplementary Figure 4), the occurrence of the T-type calcium current was prominent in DRG neurons obtained from female organ donors (Figure 7B). The T-type calcium current was recorded in 7 organ donors, 5 females and 2 males, and was absent in the remaining 3 organ donors, 2 males and 1 female (Figure 7A). On average, T-type calcium current was found in ∼27.14 % of DRG neurons obtained from female organ donors but only in 9.68 % of DRG neurons obtained from males (Figure 7B, *p*=0.0498). The total LVA calcium current amplitude measured at –40 mV and the HVA calcium current amplitude measured at 0 mV were then analyzed in all recorded neurons (n=101). The calcium current amplitude measured at –40 mV was significantly greater in DRG neurons obtained from female organ donors than in those obtained from male organ donors (Figure 6C). The calcium current density measured at –40 mV was 3.61 ± 0.75 pA/pF (n=70) and 0.74 ± 0.36 pA/pF (n=31) in DRG neurons from female and male organ donors, respectively (*p*=0.0026). In contrast, the calcium current density measured at 0 mV did not vary significantly according to the sex of the organ donor (Figure 6D).

**Figure 7.**
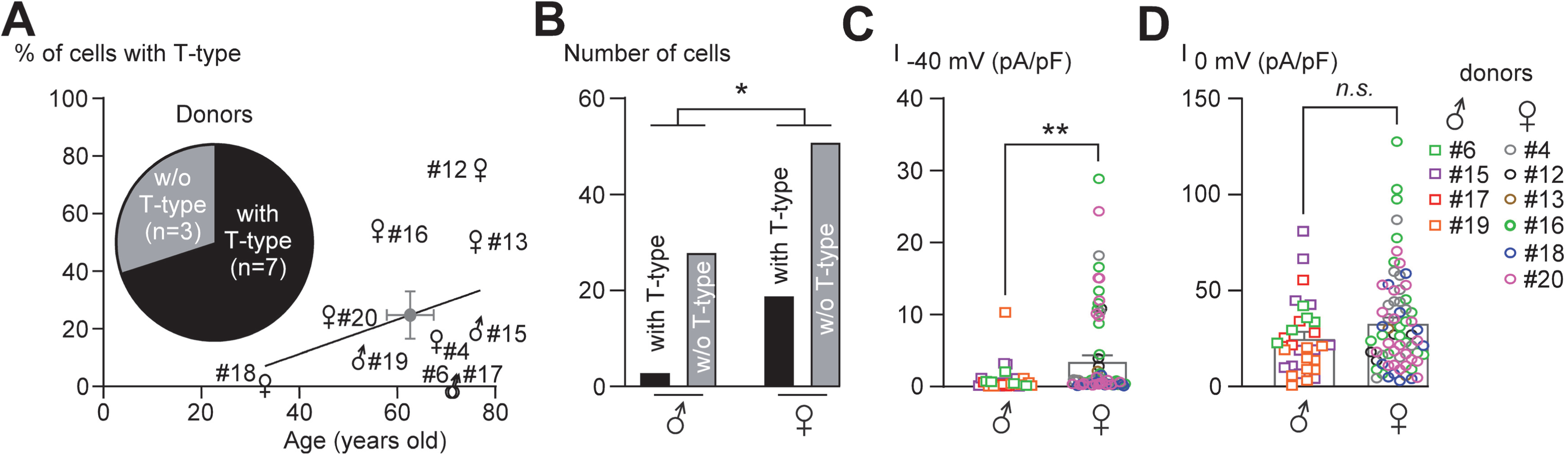
Expression of calcium currents according to the sex of organ donor and the vertebral location of DRGs. (**A**) Expression of the T-type calcium current in each organ donor according to their age and their sex (4 males and 6 females). The T-type calcium current was recorded in ∼70 % of organ donors (inset). Average age and T-type calcium current mean expression are shown in the gray circle. Non-parametric Spearman’s test and simple linear regression analysis revealed a non-significant correlation between T-type calcium current expression and age of the organ donors (all donors: *p*=0.266, *r*=0.387, N=10; male donors: *p*=0.666, *r*=0.316, N=4; and female donors: *p*=0.136, *r*=0.714, N=6). **(B)** Expression of T-type calcium current in neurons from male (n=31) and female (n=70) organ donors (contingency analysis with two-tailed chi-squared test, *p*=0.0498). **(C and D)** LVA current density (C, *p*=0.0058, n=101) and HVA current density (D, *p*=0.1752, n=101) in neurons from male and female organ donors (non-parametric two-tailed Mann-Whitney test).

## Discussion

In this study, we show that human DRG from organ donors express a functional T-type calcium channel that is primarily dependent on Cav3.2 expression. Both RNA in-situ hybridization and electrophysiological data indicate that Cav3.2 is present in a subset of human DRG neurons, representing approximately 20 % of the total neurons. Importantly, the T-type / Cav3.2 current was prominent in DRG neurons from female donors. The T-type current exhibited biophysical and pharmacological properties typical of the Cav3.2 current identified in rodent DRG neurons, including low-voltage activation, fast inactivation, and inhibition by low concentrations of nickel and Z944 (11, 16, 31–34). Although the low-voltage-activated / T-type calcium channel was first identified in chicken and rodent DRG neurons ((35–37)), which predominantly express the nickel-sensitive Cav3.2 channel (9, 11, 38), T-type calcium current expression heterogeneous between species highlighting the importance of assessing the channel expression in human DRG. Studies in rats have shown a high expression of T-type calcium current in a subset of small and medium-sized DRG neurons and an absence of T-type calcium current in large diameter neurons (12, 13, 39–41), and genetic studies in mice have identified high Cav3.2 expression in two subtypes of low-threshold mechanoreceptors (LTMRs), the Aδ-LTMRs and the C-LTMRs (10, 14, 42–44). In these neurons, Cav3.2 current increases neuronal excitability and its inhibition results in analgesia to acute mechanical and thermal stimuli in both naive and chronic pain rodent models (10, 14, 18–20, 45). On the basis of much of this work, Cav3.2 emerged as an important pharmacological target for the treatment of pain disorders, but its expression in human DRG neurons remains undemonstrated so far. Our results clearly show that the human DRG neurons expressing the *CACNA1H* / Cav3.2 RNA strongly express the TrkB receptor transcript, a marker of Aδ-LTMRs (26, 27), which in rodents also express a high density of Cav3.2 channels (10, 14, 46). In addition, human DRG neurons co-expressed the RNA for Cav3.2 and P2YR1, a marker of both Aδ−LTMRs and C-LTMRs in human and non-human primate DRG neurons (26, 27, 47), and as reported in mice (27, 43, 48, 49). In contrast, Cav3.2 is sparsely expressed in TRPV1-or Nav1.8-positive neurons, which comprise nearly all C-fibers and A∂-HTMRs. However, Cav3.2 is present in a subset of neurons that express low levels of TRPM8 (TRPM8-low). While TRPM8-high are classical cold sensors (46, 50), TRPM8-low may correspond to mechano-cold neurons as recently described in single cell transcriptomic studies (27), or to a subset of hair pull fast nociceptors that express TRPM8 and KIT (51). Compared to mouse data (14, 43, 48, 49), the question of Cav3.2 expression in putative human C-LTMR remains open. The lack of a clear molecular signature for this/these DRG subtype(s) in humans makes it difficult to answer this question properly and we could not exclude that the small fraction of Cav3.2/Nav1.8 positive cells might include these neurons. While we did not find significant RNA expression for *CACNA1I* / Cav3.3, we observed RNA expression of *CACNA1G* / Cav3.1 in Nav1.8– and TRPV1-positive-neurons. However, we found a nuclear localization of *CACNA1G* RNA puncta, likely indicating that in human DRG neurons, *CACNA1G* RNA is not fully matured and exported to the cytoplasm, resulting in the absence of functional protein (52, 53). This expression is intriguing and deserves further study to understand how the Cav3.1 RNA is regulated and possibly translated under specific conditions. Overall, our RNAscope study is in good agreement with the data obtained in rodents and the recent transcriptomic atlas of human DRG neurons indicating that Cav3.2 is the major T-type calcium channel isoform expressed primarily in Aδ−LTMRs and in a subtype of “Trpm8-low” putative cold/mechano-receptors (26, 27, 54). Future functional studies investigating the response of identified Cav3.2-positive neurons to mechanical and thermal stimuli would help to better define their properties.

A recent electrophysiological study described calcium currents in DRG neurons obtained from human donors using a method similar to ours (21). We found similar results regarding the amplitude, density, and biophysical properties of the HVA component of the calcium currents. The HVA calcium current was robustly recorded in all neurons and increased with the cell diameter and capacitance. However, our results contrast sharply with the study by Hartung et al. regarding the characteristics of the LVA current. In this previous study, the LVA current had a low density and exhibited a fast activation and very slow inactivation kinetics, atypical for the classical T-type calcium current recorded in rodent sensory neurons. The pharmacological properties of this LVA current were not investigated in Hartung et al. whereas we describe in this study that the T-type current has the expected pharmacological profile such as sensitivity to Z944 and to a low concentration of nickel. Interestingly, in a very small fraction of the neurons analyzed (∼2%), we found an atypical LVA current reminiscent of those described by Hartung et al. This current was of low amplitude, rapidly activated and slowly inactivated, and was insensitive to nickel and Z944 (which also inhibited the Cav3.1 and Cav3.3 current). The molecular characterization of this LVA current would require more in-depth pharmacological investigations. The apparent reason for the inconsistency of our T-type calcium channel characterization with the above mentioned study is not clear, although in contrast to the use of NGF by Hartung et al. for their human DRG culture procedure, we do not use any growth factors in our experiments. In addition, although our human donor sample is similar with respect to donor age, most of the data from Hartung et al. were obtained in DRG neurons from male organ donors (17 out of 21 donors).

Importantly, we found that the T-type current was more frequent and larger in DRG neurons obtained from female organ donors than in males, while a similar sex bias was not observed for the HVA current. Interestingly, RNAscope analysis showed no difference in Cav3.2 RNA between male and female organ donors confirming a transcriptomic study (55). This suggests that the increase in LVA current may involve sex-dependent post-translational modifications that strongly regulate the Cav3.2 current, such as phosphorylation (56), trafficking (57), glycosylation (17, 58, 59), or direct inhibition by neuroactive steroids and bio-active lipids (60, 61), all of which are involved in pain perception (19, 62).

Among the few analgesic molecules described as T-type calcium channel specific (63, 64), ABT-639 (22) and Z944 (30) have been evaluated in clinical trials for their ability to relieve pain in humans. Although ABT-639 was well tolerated, it did not produce analgesic effects over placebo in patients with diabetic peripheral neuropathic pain (23–25). These studies have clearly dampened the interest in considering T-type calcium channels as valuable clinical targets for innovative analgesics. However, in this study we show that ABT-639 has no significant inhibitory effect on T-type calcium currents in human DRG neurons, and we further confirmed this finding using recombinant T-type calcium Cav3.2 currents. This is consistent with recent results obtained in rodent neurons (65). This stunning lack of efficacy of ABT-639 in human DRG neurons, re-emphasizes the notion of the T-type calcium channel as a valuable target for novel analgesics for human pain relief. Interestingly, we found that Z944 is a potent inhibitor of the T-type calcium current in human DRG neurons. Accordingly, a Phase 1B clinical trial has shown that oral administration of Z944 is analgesic in laser evoked potential-induced pain in human volunteers sensitized with either capsaicin or mild UV burn injury (66). In addition, our results indicate that Cav3.2 current density is sexually dimorphic, with a prominent density in DRG neurons from female organ donors compared to those of males. Therefore, this sex bias is an important criterion to be considered in future clinical trials investigating the effect of Cav3.2 blockers for the treatment of pain, which is more prevalent in females than in males (67). Finally, the recent genetic target-disease linkage of Cav3.2 mutations in trigeminal neuralgia patients, especially with a high prevalence in females, reinforces the notion of its impact in human chronic pain (68). The prominent expression of Cav3.2 in putative Aδ-LTMRs and a subset of putative cold mechanoreceptors, supports a possible mechanistic link to mechanical and cold allodynia – pain evoked by light touch and innocuous cold, which are among the most common symptoms of neuropathic pain (69). Consistent with the involvement of these neurons in mechanical and cold allodynia, targeted genetic ablation of TrkB-positive neurons or photoablation of their skin endings are both necessary and sufficient to eliminate allodynia in neuropathic pain models (70). Taken together, our results, unambiguously demonstrating the presence of functional Cav3.2 calcium channels in a subset of human DRG neurons provide critical information for targeting these ion channels for the treatment of pain.

## Methods

### Human donors

Data were collected from a total of 20 organ donors. Chromogenic RNAscope and electrophysiology experiments were performed in Montpellier (France) on hDRGs from 16 organ donors (see donor characteristics in Table 1) obtained from anonymized organ donors at the Hospital of Montpellier, confirmed neurologically deceased, according to protocols approved by the French organ transplantation agency, Agence de la Biomédecine (DC-2014-2420). Fluorescent RNAscope experiments were performed in Dallas (USA) using an independent cohort of 4 organ donors (see donor characteristics in Table 1) for whom human tissue procurement procedures were approved by the Institutional Review Boards at the University of Texas at Dallas and hDRGs were obtained from organ donors through a collaboration with the Southwest Transplant Alliance.

### DRG extraction

#### Montpellier procedure

Organ donor body temperature was lowered with ice, and blood circulation was maintained for 3 hours before the vertebral bloc removal through a ventral approach as previously described (71). A spinal segment from thoracic (T9) to lumbar (L2) was removed in 1 piece, and DRGs were immediately dissected and either 1) maintained in ice-cold, oxygenated calcium/magnesium free HBSS/Glucose/pen-strep (HBSS^G-AB^) for less than 30 minutes until returned to the laboratory for cell culture and subsequent electrophysiological analysis; or 2) flash frozen, for subsequent ISH experiments.

#### Dallas procedure

DRGs were recovered (∼2 hours post-mortem) using a ventral approach as previously described (72). L1 and L2 lumbar DRGs were collected and flash frozen for subsequent ISH experiments.

### RNAscope In Situ Hybridization

For all experiments performed, DRG tissue blocks in the OCT were removed from the –80°C freezer, placed on dry ice, and transferred to the laboratory. DRGs were sectioned at 16µm (chromogenic RNAscope) or 20μm (fluorescent RNAscope) on SuperFrost Plus loaded slides (Fisher Scientific; Cat 12-550-15) in a cryostat. Sections were thawed only briefly to adhere to the slide, but immediately returned to the –20°C cryostat chamber until sectioning was complete. Three sections (technical replicates) from 3-4 donors (biological replicates) were stained in each experiment. Slides were removed from the cryostat and immediately immersed in cold (4°C) 10% formalin (Fisher Scientific; Cat. 23-245684) for 15 minutes. The tissues were then dehydrated sequentially in 50% ethanol (5 min; Fisher Scientific; Cat 04-355-223), 70% ethanol (5 min), and twice in 100% ethanol (5 min) at room temperature. Slides were briefly air-dried, and then borders were drawn around each section using a hydrophobic pen (ImmEdge PAP pen; Vector Labs). After the hydrophobic borders dried, the slides were immediately processed for RNAscope in situ hybridization.

RNAscope in situ hybridization chromogenic duplex and fluorescent multiplex version 2 (Advanced Cell Diagnostics (ACD); Cat. 322430 and 323100, respectively) was performed on human DRGs using the fresh-frozen protocol as described by ACD (Manual #3225500-USM and 323100-USM with revision date: 04092019 and 02272019). Hydrogen peroxide (ACD; Cat. 322381) was applied to each section until completely covered and incubated for 10 minutes at room temperature. The slides were then washed in distilled water and incubated individually in protease III reagent (ACD; Cat. No. 322381) for 10 seconds at room temperature.

The protease incubation time was optimized for the tissue and specific lot of protease reagent as recommended by ACD. RNAscope assays were performed according to the manufacturer’s instructions using a HybEZ oven (ACD; Cat. 321710). The probes used were Hs-cacna1h-C1 and C2 (ACD; Cat. 528311 and 528311-C2), Hs-cacna1g-C1 (ACD; Cat. 528321), Hs-cacna1i-C1 (ACD; Cat. 459971-C1), Hs-scn10a-C1 and C2 (ACD; Cat. 406291 and 406291-C2), Hs-STT1-C2 (ACD; Cat. 525791-C2), Hs-NTRK2-C2 (ACD; 402621-C2), Hs-P2ry1-C1 (ACD; cat. 485831), Hs-Trpm8-C1 (ACD; cat. 543121), Hs-Trpv1-C2 (ACD; cat. 415381), Hs-AIF1-C3 (ACD; cat. 433121-C3), positive control Hs-UBC-POLR2A-PPIB-C123 (ACD; cat. 320861), negative control dapB-C123 (ACD; cat. 320871). For chromogenic ISH, hematoxylin counterstaining (30% Gill hematoxylin n°1; Sigma-Aldrich) was performed at the end of the procedure, and slides were mounted with Vectamount mounting medium (Vector Labs). For fluorescence ISH, TSA Plus Akoya dyes in fluorescein, cyanine-3, and cyanine-5 (Akoya; NEL741001KT, NEL744001KT, NEL745001KT) were used. Akoya cyanine-3, fluorescein, and cyanine-5 dyes were assigned to C1, C2, and C3 probes, respectively. DAPI (ACD; Cat. 323110) was applied to each section for 1 minute at room temperature at the end of the procedure, then washed, air dried, and coverslipped (Globe Scientific; Cat. 1415-15) with Prolong Gold Antifade mounting medium (Fisher Scientific; Cat. P36930).

A positive and negative control was run on a single section from each DRG for every RNAscope experiment. The positive control probe cocktail contains probes for high, medium, and low-expressing mRNAs that are present in all cells (ubiquitin C > Peptidyl-prolyl cis-trans isomerase B > DNA-directed RNA polymerase II subunit RPB1) and allows us to gauge tissue quality and experimental conditions. All tissues showed robust signal for all positive control probes (C1 and C2 for chromogenic ISH, C1, 2 and 3 for fluorescent ISH). A negative control probe against the bacterial DapB gene was used to check for lipofuscin and background labeling. Chromogenic ISH allowed separating the brown lipofuscin signal from the specific C1-blue and C2-pink signals. We carefully excluded the lipofuscin fluorescent background from the specific ISH signal with the multiplex RNAscope approach as published previously (73, 74).

For chromogenic ISH, slides were imaged at 40x with Hamamastu Nanozoomer 2 slide scanner. The scanner was configured to take a Z-stack of three images of each DRG section with a focus adjustment. Slides images were viewed with the NDP-view2 (Hamamatsu; V 2.9.29) software to take individual tiff images of each DRG at maximum resolution. These images were then analyzed with Fiji freeware using the cell counter plugin, and the measure tool to determine DRG cross section area. Fluorescent ISH, slides were imaged on a FV3000 confocal microscope (Evident Scientific) at 10X or 20X magnification. For quantification, three 20X images were acquired from each DRG tissue section, and three sections were imaged per DRG for a total of 9 images. Acquisition parameters were set based on guidelines for the FV3000 provided by Evident Scientific. The raw image files were analyzed in CellSens (Olympus; v1.18). Lipofuscin quenchers (such as True Black) are not compatible with RNAscope. Large globular structures and/or signal that autofluoresced in all channels (particularly brightest at 488 and 555 wavelengths) were considered to be background lipofuscin and were not analyzed. Aside from adjusting brightness/contrast, we performed no digital image processing to subtract background. For both chromogenic and fluorescent images, DRG were considered positive for a labeled mRNA if they had greater than 5 positive puncta. The diameter of Cav3.1, Cav3.2, and Cav3.3 positive neurons with a visible nucleus was measured using the polyline tool in CellSens. Neurons were also tracked for enrichment of Cav3.1, Cav3.2, and Cav3.3 mRNA molecules using two different criteria: >50 puncta (robust) and <50 puncta (low), although within the low category most of the neurons fell within the 6-35 puncta range. Since the percentage of TRPV1+ neurons was calculated in the Cav3.1, Cav3.2, and Cav3.3 experiments from the same donor and the same DRG, the values for each DRG was averaged across all three experiments. Graphs were generated using GraphPad Prism version 10 (GraphPad Software, Inc. San Diego, CA USA).

### Isolation and plating of human dorsal root ganglion neurons

Electrophysiological experiments were performed on isolated lumbar and thoracic dorsal root ganglion (DRG) neurons from 10 organ donors (6 females and 4 males, aged 33 to 77 years (62.6 ± 4.7 years old)). Within 30 to 60 minutes after dissection, two to three 3 DRGs per donor were cleaned and minced into small pieces and incubated at 37°C in 8ml of HBSS^G-AB^ solution containing 4 mg/ml collagenase, 10mg/ml dispase (Sigma Aldrich, France) and 5mM calcium with gentle shaking. Tissue digestion was estimated by observing the release of isolated DRG somata after triturating a piece of tissue with a fire-polished, large-aperture Pasteur glass pipette. Tissues were further incubated for an additional 30 to 90 minutes as needed. Digestion was stopped by washing the tissues twice with 4 ml HBSS^G-AB^ without calcium, and 2 times with culture medium (Neurobasal A / B27 / L-glutamine / pen-strep). Trituration was performed with Pasteur pipettes of decreased tips diameters. Debris were allowed to settle down the tube for 2 minutes and cell suspension was collected, passed through a 100µm cell strainer (Ducher France; clearline 141380C), and centrifuged at 500g for 5 minutes. Cells were suspended in culture media and seeded on plastic Petri dishes coated with poly D-Lysine and laminin A and kept at 37°C in an humidified culture incubator. The culture media was not supplemented with growth factors. Cells were used for patch-clamp experiments from 6 to 72 hours after cell preparation with most recordings made within 36 hours.

### Electrophysiology

Patch clamp recordings were performed using an Axopatch 200B amplifier (Molecular Devices, Sunnyvale CA) and fire-polished borosilicate glass pipettes with a typical resistance of 1.5–2.5 MOhm. Macroscopic calcium currents were recorded at room temperature using an internal solution containing (in mM): 100 CsCl, 40 tetraethylammonium (TEA)-Cl, 10 EGTA, 10 HEPES, 3 Mg-ATP, 0.6 GTPNa, and 3 CaCl2 (pH adjusted to 7.25 with KOH, ∼300 mOsm, ∼100 nM free Ca2+ calculated with the MaxChelator software) and an extracellular solution containing (in mM): 140 TEA-Cl, 10 4-Aminopyridin, 5 KCl, 2 CaCl2, 2 NaCl, 1 MgCl2, 10 HEPES and 10 Glucose (pH adjusted to 7.25 with TEA-OH, ∼310 mOsm). Extracellular solutions were applied by a gravity-driven homemade perfusion. Recordings were filtered at 5 kHz. In the double pulse protocol, a P/8 leak subtraction was performed. Data were analyzed using pCLAMP10 (Molecular Devices) and GraphPad Prism 9 (GraphPad) software. To account for the presence of a low-voltage activated (LVA) and a high-voltage activated (HVA) calcium current, current-voltage (IV) curves were fitted using a double combined Boltzmann and linear ohmic relationships, where *I=*(Gmax_(LVA)_*(Vm-Erev_(LVA)_)/(1+exp((Vm-V0.5_(LVA)_)/slope_(LVA)_)))+(Gmax_(HVA)_*(Vm-Erev_(HVA)_)/(1+exp((Vm-V0.5_(HVA)_)/slope_(HVA)_))).To minimize the consequence of current rectification near reversal potential on the determination of conductance, the current values greater than +30 mV were not considered for the fit. The normalized conductance-voltage curve for LVA current was then constructed with a Boltzmann equation: *G*/*G*max = 1/(1 + *exp*(V0.5_(LVA)_-Vm)/slope_(LVA)_). Steady-state inactivation curves were fitted using the Boltzmann equation where I/I max = 1/(1+exp((Vm-V0.5)/slope factor)). Time constants (Tau) of activation and inactivation were extracted from double-exponential fit of the current traces at –40 mV (LVA current) and 0 mV (HVA current) and deactivation time constant were extracted from mono-exponential fit of the current traces recorded at –90 mV. Electrophysiological procedures on HEK-293 cells were performed as previously described (75). Results are presented as the mean ± SEM, and n is the number of cells whereas N is the number of organ donors. Statistical analysis was performed as indicated in the Figure legends (* *p*<0.05, ** *p*<0.01, *** *p*<0.001, **** *p*<0.0001).

### Chemical reagents

Compounds were purchased from Sigma Aldrich (Merck France), excepted for ABT-639 that was purchased from Tocris (Bio-Techn, France). Z944 is a generous gift of Dr T.P. Snutch.(University of British Columbia, Vancouver, Canada).

## Author contributions

JC, PFM, TJP, EB designed experiments, JC, VS, SS, AF, GP, NL, FV, LB, PFM, EB conducted the experiments, JC, VS, SS, PFM, TJP, EB analyzed the data, and JC TJP and EB wrote the manuscript.

## Acknowledgements

We gratefully acknowledge the gift of human tissue from all donors included in this work and their families, whose contribution has been critical for this work. This work was supported by ANR-15-CE16-0012-PainT, ANR-21-CE44-0006-interacT, Labex-ICST-11-LABX-0015, FRC-EET-2019, FHU-inovPain2.0 grants (EB); by CNRS/INSERM/Montpellier Hospital research supports; and by NIH grant NS130608 (TJP).

## Supplementary Figure Legends

**Supplementary Figure 1.**
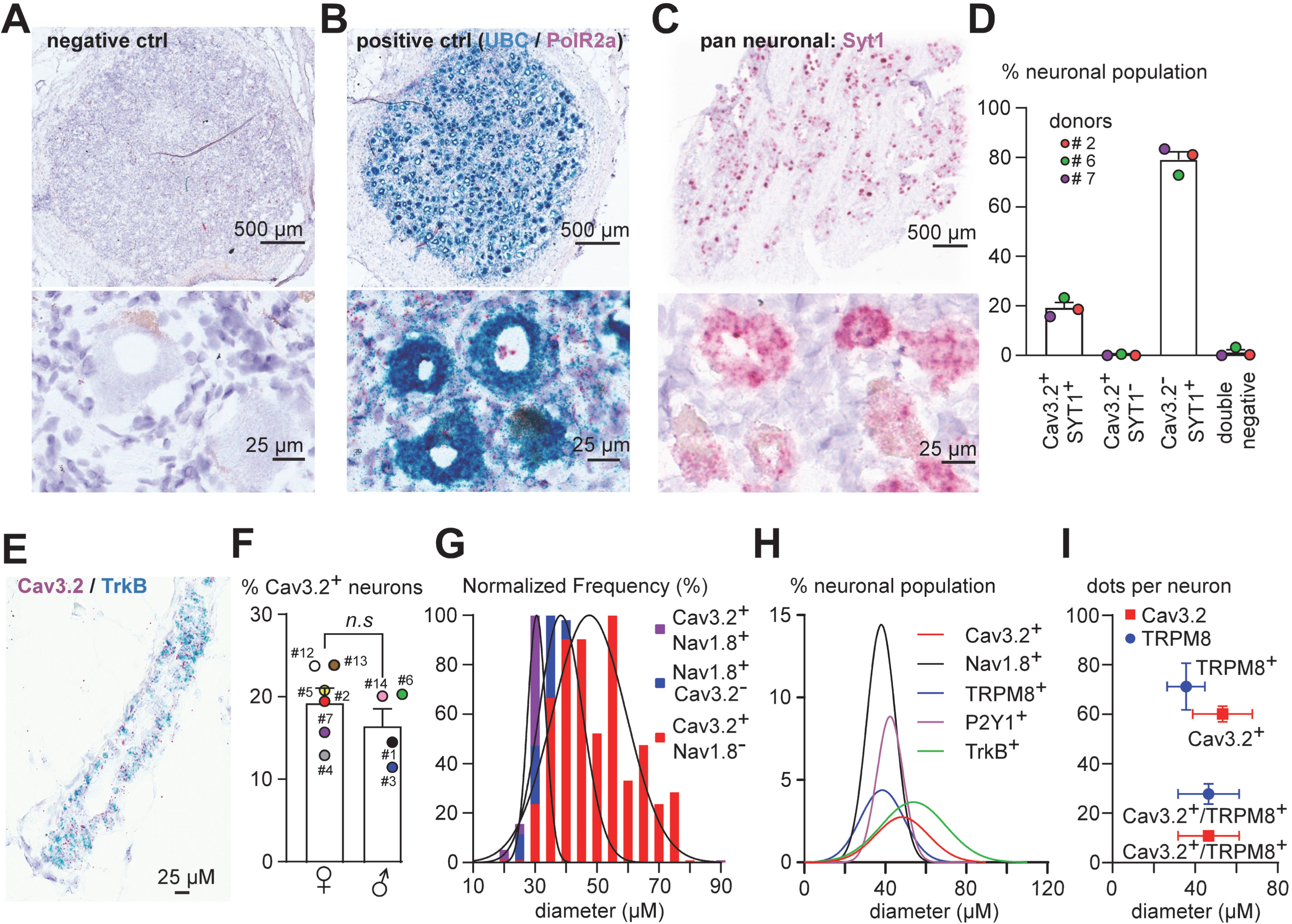
RNAscope experiments in human dorsal root ganglia. (**A-C**) Representative image of hDRGs labeled with RNAscope chromogenic *in situ* hybridization for DapB (A, negative control), UBC (B, positive control, highly expressed RNA), PolR2a (B, positive control, weakly expressed RNA), and Syt1 (C, a pan-neuronal marker). **(D)** Coexpression of Cav3.2 and Syt1 in hDRG neurons. **(E)** Expression of Cav3.2 and TrkB in a DRG blood vessel. **(F)** Cav3.2 expression in DRG neurons from female and male organ donors (non-parametric two-tailed Mann-Whitney test, *p*=0.352). **(G)** Normalized frequency (%) of hDRG neurons expressing Cav3.2 and/or Nav1.8 relative to cell diameter. **(H)** Relative frequency (% of total neuronal population) of hDRG neurons expressing Cav3.2, Nav1.8, TRPM8, P2Y1, TrkB and TRPV1 relative to cell diameter. **(I)** Quantitative expression (number of RNA dots) of Cav3.2 (red) and TRPM8 (blue) in Cav3.2^+^/TRPM8^-^ neurons (Cav3.2^+^) or Cav3.2^-^/TRPM8^+^ neurons (TRPM8^+^) compared to Cav3.2^+^/TRPM8^+^ neurons (non-parametric two-tailed Mann-Whitney test, *p*<0.0001).

**Supplementary Figure 2.**
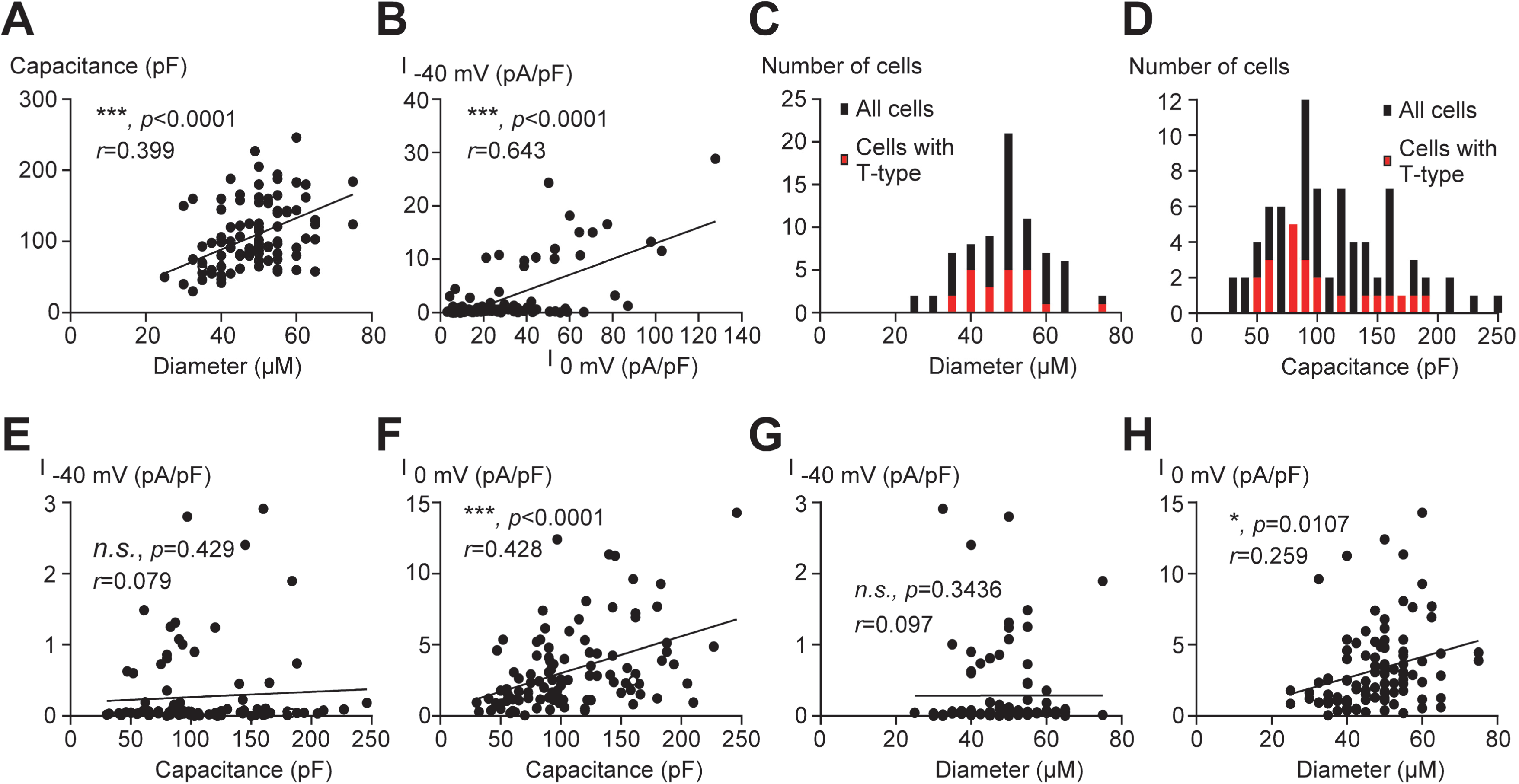
Expression of calcium current in relation to cell diameter and membrane capacitance. (**A**). Positive correlation between cell diameter and membrane capacitance (non-parametric Spearman’s test and simple linear regression analysis, n=101). **(B).** Positive correlation between calcium current density measured at –40 mV (I _-40 mV_) and at 0 mV (I _0 mV_, non-parametric Spearman’s test and simple linear regression analysis, n=101). **(C and D)** Frequency distribution of the number of cells expressing or not expressing the T-type calcium current as a function of cell diameter (C, n=101) and membrane capacitance (D, n=101). **(E and F)** Correlation between cell membrane capacitance and calcium current amplitude measured at – 40 mV (E, I _-40 mV_) or at 0 mV (F, I _0 mV_, non-parametric Spearman’s test and simple linear regression analysis, n=101). **(G and H)** Correlation between cell diameter and calcium current amplitude measured at –40 mV (G, I _-40 mV_) or at 0 mV (H, I _0 mV_, non-parametric Spearman’s test and simple linear regression analysis, n=101).

**Supplementary Figure 3.**
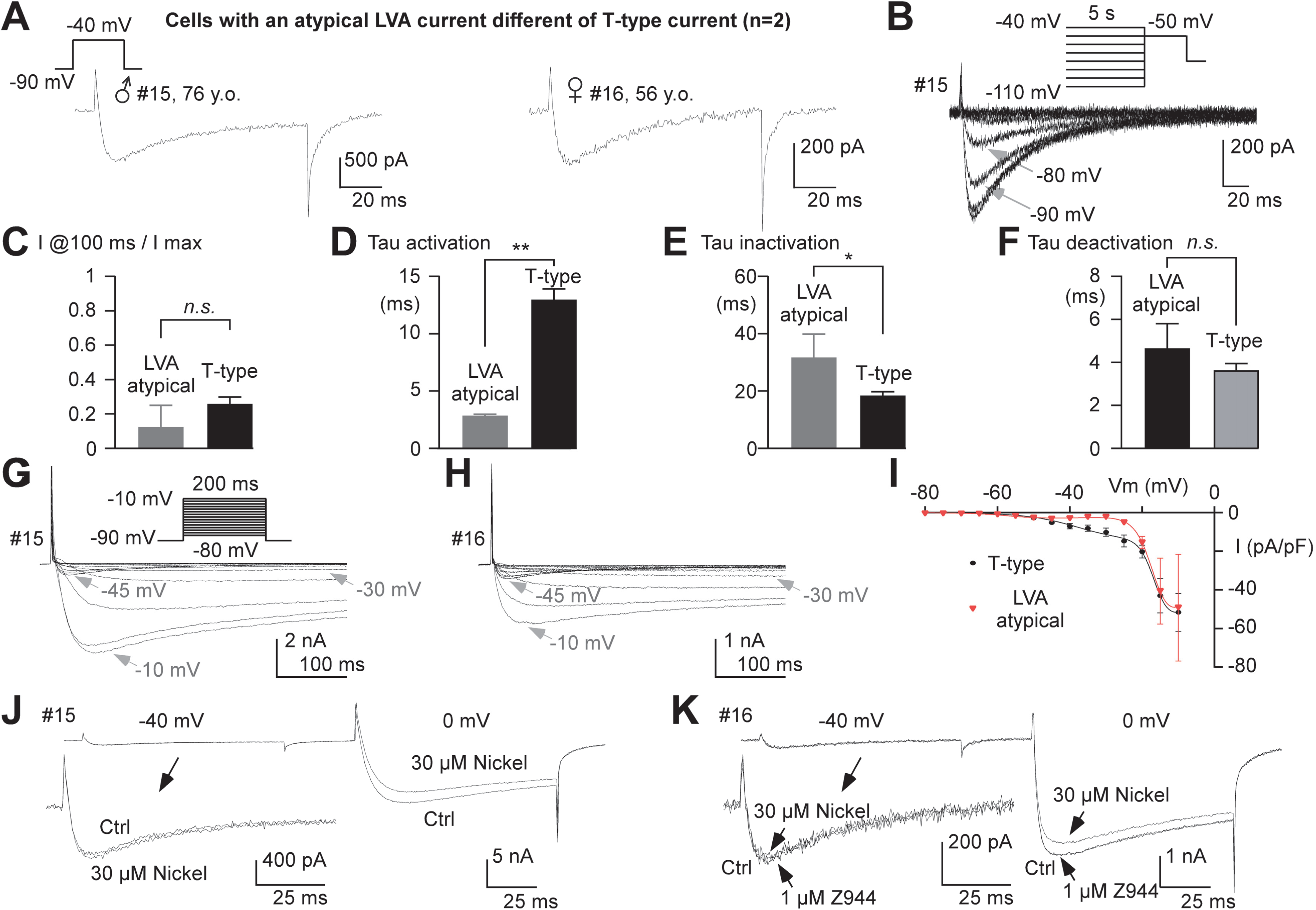
Recordings in 2 neurons expressing an atypical LVA current with a pharmacology distinct from the T-type calcium current. (**A**) Atypical LVA current recorded in organ donors #15 and #16. **(B)** Steady-state inactivation of the LVA current recorded in organ donor #16. **(C)** Mean normalized current amplitude at 100 ms of the atypical LVA current (n=2) compared to the T-type calcium current (*n.s.*, non-parametric two-tailed Mann-Whitney test, *p*=0.3116, n=22). **(D)** Activation kinetics of the atypical LVA current (n=2) compared to the T-type calcium current (\**\**, non-parametric two-tailed Mann-Whitney test, *p*=0.0072, n=22). **(E)** Inactivation kinetics of the atypical LVA current (n=2) compared to the T-type calcium current (*\**, non-parametric two-tailed Mann-Whitney test, *p*=0.0435, n=22). **(F)** Deactivation kinetics of the atypical LVA current (n=2) compared to the T-type calcium current (*n.s.*, non-parametric two-tailed Mann-Whitney test, *p*=0.3874, n=22). **(G-H)** Calcium currents elicited by a series of step depolarizations ranging from −80 to +40 mV (0.2 Hz stimulation frequency) from a holding potential (HP) of −90 mV in organ donor #15 (G) and #16 (H). Note that the atypical LVA current peaked at –45 mV. **(I)** Current-voltage (I-V) curves obtained in cells expressing the atypical LVA current (n=2, red) and in cells expressing the T-type calcium current (n=16, black). **(J)** Effect of 30 µM nickel on the atypical LVA current recorded at –40 mV and on the HVA calcium current recorded at 0 mV in organ donor #15. **(K)** Effect of 30 µM nickel and of 1 µM Z944 on the atypical LVA current recorded at –40 mV and on the HVA calcium current recorded at 0 mV in organ donor #16.

**Supplementary Figure 4.**
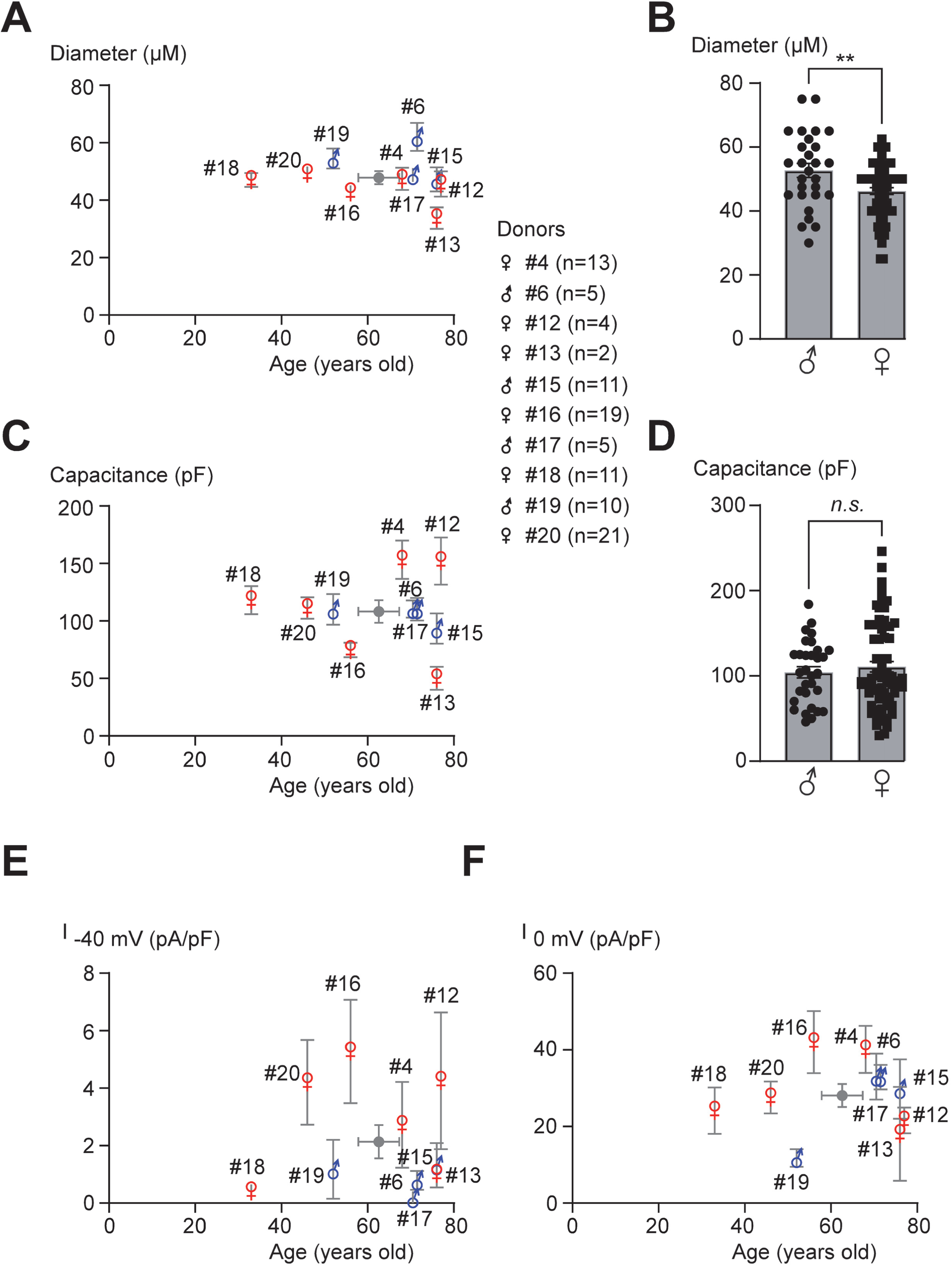
Calcium current densities as a function of sex and age of organ donors. (**A**) Mean calcium current density measured at –40 mV (I _-40 mV_) in each organ donor according to their sex and age. Mean age and calcium current density are shown in the gray circle. Non-parametric Spearman test and simple linear regression analysis revealed non-significant correlation between current density and organ donor age (*p*=0.804, *r*=0.0912, N=10 for all donors). **(B)** Mean calcium current density measured at 0 mV (I _0 mV_) in each organ donor according to their sex and age. Mean age and calcium current density are shown in the gray circle. Non-parametric Spearman test and simple linear regression analysis revealed non-significant correlation between current density and organ donor age (*p*=0.857, *r*=-0.0668, N=10).

## References

1. Cohen SP, Vase L, and Hooten WM. Chronic pain: an update on burden, best practices, and new advances. Lancet. 2021;397(10289):2082–97.

2. Vowles KE, McEntee ML, Julnes PS, Frohe T, Ney JP, and van der Goes DN. Rates of opioid misuse, abuse, and addiction in chronic pain: a systematic review and data synthesis. Pain. 2015;156(4):569–76.

3. Renthal W, Chamessian A, Curatolo M, Davidson S, Burton M, Dib-Hajj S, et al. Human cells and networks of pain: Transforming pain target identification and therapeutic development. Neuron. 2021;109(9):1426–9.

4. Middleton SJ, Barry AM, Comini M, Li Y, Ray PR, Shiers S, et al. Studying human nociceptors: from fundamentals to clinic. Brain. 2021;144(5):1312–35.

5. Waxman SG, and Zamponi GW. Regulating excitability of peripheral afferents: emerging ion channel targets. Nat Neurosci. 2014;17(2):153–63.

6. Zamponi GW, Striessnig J, Koschak A, and Dolphin AC. The Physiology, Pathology, and Pharmacology of Voltage-Gated Calcium Channels and Their Future Therapeutic Potential. Pharmacol Rev. 2015;67(4):821–70.

7. Ertel EA, Campbell KP, Harpold MM, Hofmann F, Mori Y, Perez-Reyes E, et al. Nomenclature of voltage-gated calcium channels. Neuron. 2000;25(3):533–5.

8. Chemin J, Monteil A, Perez-Reyes E, Bourinet E, Nargeot J, and Lory P. Specific contribution of human T-type calcium channel isotypes (alpha(1G), alpha(1H) and alpha(1I)) to neuronal excitability. J Physiol. 2002;540(Pt 1):3–14.

9. Perez-Reyes E. Molecular physiology of low-voltage-activated t-type calcium channels. Physiol Rev. 2003;83(1):117–61.

10. Shin JB, Martinez-Salgado C, Heppenstall PA, and Lewin GR. A T-type calcium channel required for normal function of a mammalian mechanoreceptor. Nat Neurosci. 2003;6(7):724–30.

11. Bourinet E, Alloui A, Monteil A, Barrere C, Couette B, Poirot O, et al. Silencing of the Cav3.2 T-type calcium channel gene in sensory neurons demonstrates its major role in nociception. EMBO J. 2005;24(2):315–24.

12. Nelson MT, Joksovic PM, Perez-Reyes E, and Todorovic SM. The endogenous redox agent L-cysteine induces T-type Ca2+ channel-dependent sensitization of a novel subpopulation of rat peripheral nociceptors. J Neurosci. 2005;25(38):8766–75.

13. Coste B, Crest M, and Delmas P. Pharmacological dissection and distribution of NaN/Nav1.9, T-type Ca2+ currents, and mechanically activated cation currents in different populations of DRG neurons. J Gen Physiol. 2007;129(1):57–77.

14. Francois A, Schuetter N, Laffray S, Sanguesa J, Pizzoccaro A, Dubel S, et al. The Low-Threshold Calcium Channel Cav3.2 Determines Low-Threshold Mechanoreceptor Function. Cell Rep. 2015;10(3):370–82.

15. Messinger RB, Naik AK, Jagodic MM, Nelson MT, Lee WY, Choe WJ, et al. In vivo silencing of the Ca(V)3.2 T-type calcium channels in sensory neurons alleviates hyperalgesia in rats with streptozocin-induced diabetic neuropathy. Pain. 2009;145(1-2):184–95.

16. Marger F, Gelot A, Alloui A, Matricon J, Ferrer JF, Barrere C, et al. T-type calcium channels contribute to colonic hypersensitivity in a rat model of irritable bowel syndrome. Proc Natl Acad Sci U S A. 2011;108(27):11268–73.

17. Orestes P, Osuru HP, McIntire WE, Jacus MO, Salajegheh R, Jagodic MM, et al. Reversal of neuropathic pain in diabetes by targeting glycosylation of Ca(V)3.2 T-type calcium channels. Diabetes. 2013;62(11):3828–38.

18. Todorovic SM, and Jevtovic-Todorovic V. Neuropathic pain: role for presynaptic T-type channels in nociceptive signaling. Pflugers Arch. 2013;465(7):921–7.

19. Cai S, Gomez K, Moutal A, and Khanna R. Targeting T-type/CaV3.2 channels for chronic pain. Transl Res. 2021;234:20–30.

20. Harding EK, and Zamponi GW. Central and peripheral contributions of T-type calcium channels in pain. Mol Brain. 2022;15(1):39.

21. Hartung JE, Moy JK, Loeza-Alcocer E, Nagarajan V, Jostock R, Christoph T, et al. Voltage-gated calcium currents in human dorsal root ganglion neurons. Pain. 2022;163(6):e774–e85.

22. Jarvis MF, Scott VE, McGaraughty S, Chu KL, Xu J, Niforatos W, et al. A peripherally acting, selective T-type calcium channel blocker, ABT-639, effectively reduces nociceptive and neuropathic pain in rats. Biochem Pharmacol. 2014;89(4):536–44.

23. Serra J, Duan WR, Locke C, Sola R, Liu W, and Nothaft W. Effects of a T-type calcium channel blocker, ABT-639, on spontaneous activity in C-nociceptors in patients with painful diabetic neuropathy: a randomized controlled trial. Pain. 2015;156(11):2175–83.

24. Ziegler D, Duan WR, An G, Thomas JW, and Nothaft W. A randomized double-blind, placebo-, and active-controlled study of T-type calcium channel blocker ABT-639 in patients with diabetic peripheral neuropathic pain. Pain. 2015;156(10):2013–20.

25. Wallace M, Duan R, Liu W, Locke C, and Nothaft W. A Randomized, Double-Blind, Placebo-Controlled, Crossover Study of the T-Type Calcium Channel Blocker ABT-639 in an Intradermal Capsaicin Experimental Pain Model in Healthy Adults. Pain Med. 2016;17(3):551–60.

26. Jung M, Dourado M, Maksymetz J, Jacobson A, Laufer BI, Baca M, et al. Cross-species transcriptomic atlas of dorsal root ganglia reveals species-specific programs for sensory function. Nat Commun. 2023;14(1):366.

27. Bhuiyan SA, Xu M, Yang L, Semizoglou E, Bhatia P, Pantaleo KI, et al. Harmonized cross-species cell atlases of trigeminal and dorsal root ganglia. Sci Adv. 2024;10(25):eadj9173.

28. Yaari Y, Hamon B, and Lux HD. Development of two types of calcium channels in cultured mammalian hippocampal neurons. Science. 1987;235(4789):680–2.

29. Lee JH, Gomora JC, Cribbs LL, and Perez-Reyes E. Nickel block of three cloned T-type calcium channels: low concentrations selectively block alpha1H. Biophys J. 1999;77(6):3034–42.

30. Tringham E, Powell KL, Cain SM, Kuplast K, Mezeyova J, Weerapura M, et al. T-type calcium channel blockers that attenuate thalamic burst firing and suppress absence seizures. Sci Transl Med. 2012;4(121):121ra19.

31. Todorovic SM, and Lingle CJ. Pharmacological properties of T-type Ca2+ current in adult rat sensory neurons: effects of anticonvulsant and anesthetic agents. J Neurophysiol. 1998;79(1):240–52.

32. Jagodic MM, Pathirathna S, Nelson MT, Mancuso S, Joksovic PM, Rosenberg ER, et al. Cell-specific alterations of T-type calcium current in painful diabetic neuropathy enhance excitability of sensory neurons. J Neurosci. 2007;27(12):3305–16.

33. Francois A, Kerckhove N, Meleine M, Alloui A, Barrere C, Gelot A, et al. State-dependent properties of a new T-type calcium channel blocker enhance Ca(V)3.2 selectivity and support analgesic effects. Pain. 2013;154(2):283–93.

34. Gong Y, Liu R, Zha H, Dong D, Lu N, Yan H et al. Analgesic Buxus alkaloids with Enhanced Selectivity for the Low-Voltage-Gated Calcium Channel Ca(v) 3.2 over Ca(v) 34.1 through a New Binding Mode. Angew Chem Int Ed Engl. 2024;63(1):e202313461.

35. Carbone E, and Lux HD. A low voltage-activated, fully inactivating Ca channel in vertebrate sensory neurones. Nature. 1984;310(5977):501-2.

36. Nowycky MC, Fox AP, and Tsien RW. Three types of neuronal calcium channel with different calcium agonist sensitivity. Nature. 1985;316(6027):440-3.

37. Bossu JL, Feltz A, and Thomann JM. Depolarization elicits two distinct calcium currents in vertebrate sensory neurones. Pflugers Arch. 1985;403(4):360–8.

38. Talley EM, Cribbs LL, Lee JH, Daud A, Perez-Reyes E, and Bayliss DA. Differential distribution of three members of a gene family encoding low voltage-activated (T-type) calcium channels. J Neurosci. 1999;19(6):1895–911.

39. White G, Lovinger DM, and Weight FF. Transient low-threshold Ca2+ current triggers burst firing through an afterdepolarizing potential in an adult mammalian neuron. Proc Natl Acad Sci U S A. 1989;86(17):6802–6.

40. Schroeder JE, Fischbach PS, and McCleskey EW. T-type calcium channels: heterogeneous expression in rat sensory neurons and selective modulation by phorbol esters. J Neurosci. 1990;10(3):947–51.

41. Scroggs RS, and Fox AP. Calcium current variation between acutely isolated adult rat dorsal root ganglion neurons of different size. J Physiol. 1992;445:639–58.

42. Dubreuil AS, Boukhaddaoui H, Desmadryl G, Martinez-Salgado C, Moshourab R, Lewin GR, et al. Role of T-type calcium current in identified D-hair mechanoreceptor neurons studied in vitro. J Neurosci. 2004;24(39):8480–4.

43. Reynders A, Mantilleri A, Malapert P, Rialle S, Nidelet S, Laffray S, et al. Transcriptional Profiling of Cutaneous MRGPRD Free Nerve Endings and C-LTMRs. Cell Rep. 2015;10(6):1007–19.

44. Bernal Sierra YA, Haseleu J, Kozlenkov A, Begay V, and Lewin GR. Genetic Tracing of Ca(v)3.2 T-Type Calcium Channel Expression in the Peripheral Nervous System. Front Mol Neurosci. 2017;10:70.

45. Wang R, and Lewin GR. The Cav3.2 T-type calcium channel regulates temporal coding in mouse mechanoreceptors. J Physiol. 2011;589(Pt 9):2229–43.

46. Qi L, Iskols M, Shi D, Reddy P, Walker C, Lezgiyeva K, et al. A mouse DRG genetic toolkit reveals morphological and physiological diversity of somatosensory neuron subtypes. Cell. 2024;187(6):1508–26 e16.

47. Kupari J, Usoskin D, Parisien M, Lou D, Hu Y, Fatt M, et al. Single cell transcriptomics of primate sensory neurons identifies cell types associated with chronic pain. Nat Commun. 2021;12(1):1510.

48. Sharma N, Flaherty K, Lezgiyeva K, Wagner DE, Klein AM, and Ginty DD. The emergence of transcriptional identity in somatosensory neurons. Nature. 2020;577(7790):392-8.

49. Huzard D, Martin M, Maingret F, Chemin J, Jeanneteau F, Mery PF, et al. The impact of C-tactile low-threshold mechanoreceptors on affective touch and social interactions in mice. Sci Adv. 2022;8(26):eabo7566.

50. Bautista DM, Siemens J, Glazer JM, Tsuruda PR, Basbaum AI, Stucky CL, et al. The menthol receptor TRPM8 is the principal detector of environmental cold. Nature. 2007;448(7150):204-8.

51. Bouchatta O, Brodzki M, Manouze H, Carballo GB, Kindstrom E, de-Faria FM, et al. PIEZO2-dependent rapid pain system in humans and mice. bioRxiv. 2023.

52. Luo MJ, and Reed R. Splicing is required for rapid and efficient mRNA export in metazoans. Proc Natl Acad Sci U S A. 1999;96(26):14937–42.

53. Garland W, and Jensen TH. Nuclear sorting of RNA. Wires Rna. 2020;11(2).

54. Yu H, Nagi SS, Usoskin D, Hu Y, Kupari J, Bouchatta O, et al. Leveraging deep single-soma RNA sequencing to explore the neural basis of human somatosensation. Nat Neurosci. 2024;27(12):2326–40.

55. North RY, Li Y, Ray P, Rhines LD, Tatsui CE, Rao G, et al. Electrophysiological and transcriptomic correlates of neuropathic pain in human dorsal root ganglion neurons. Brain. 2019;142(5):1215–26.

56. Blesneac I, Chemin J, Bidaud I, Huc-Brandt S, Vandermoere F, and Lory P. Phosphorylation of the Cav3.2 T-type calcium channel directly regulates its gating properties. Proc Natl Acad Sci U S A. 2015;112(44):13705–10.

57. Garcia-Caballero A, Gadotti VM, Stemkowski P, Weiss N, Souza IA, Hodgkinson V, et al. The deubiquitinating enzyme USP5 modulates neuropathic and inflammatory pain by enhancing Cav3.2 channel activity. Neuron. 2014;83(5):1144–58.

58. Latham JR, Pathirathna S, Jagodic MM, Choe WJ, Levin ME, Nelson MT, et al. Selective T-type calcium channel blockade alleviates hyperalgesia in ob/ob mice. Diabetes. 2009;58(11):2656–65.

59. Weiss N, Black SA, Bladen C, Chen L, and Zamponi GW. Surface expression and function of Cav3.2 T-type calcium channels are controlled by asparagine-linked glycosylation. Pflugers Arch. 2013;465(8):1159–70.

60. Pathirathna S, Todorovic SM, Covey DF, and Jevtovic-Todorovic V. 5alpha-reduced neuroactive steroids alleviate thermal and mechanical hyperalgesia in rats with neuropathic pain. Pain. 2005;117(3):326–39.

61. Chemin J, Cazade M, and Lory P. Modulation of T-type calcium channels by bioactive lipids. Pflug Arch Eur J Phy. 2014;466(4):689–700.

62. Todorovic SM, and Jevtovic-Todorovic V. T-type voltage-gated calcium channels as targets for the development of novel pain therapies. Br J Pharmacol. 2011;163(3):484–95.

63. Nam G. T-type calcium channel blockers: a patent review (2012-2018). Expert Opin Ther Pat. 2018;28(12):883–901.

64. Weiss N, and Zamponi GW. T-Type Channel Druggability at a Crossroads. ACS Chem Neurosci. 2019;10(3):1124–6.

65. Antunes FTT, Huang S, Chen L, and Zamponi GW. Effect of ABT-639 on Cav3.2 channel activity and its analgesic actions in mouse models of inflammatory and neuropathic pain. Eur J Pharmacol. 2024;967:176416.

66. Lee M. Z944: a first in class T-type calcium channel modulator for the treatment of pain. J Peripher Nerv Syst. 2014;19 Suppl 2:S11–2.

67. Johnston KJA, Ward J, Ray PR, Adams MJ, McIntosh AM, Smith BH, et al. Sex-stratified genome-wide association study of multisite chronic pain in UK Biobank. PLoS Genet. 2021;17(4):e1009428.

68. Dong W, Jin SC, Allocco A, Zeng X, Sheth AH, Panchagnula S, et al. Exome Sequencing Implicates Impaired GABA Signaling and Neuronal Ion Transport in Trigeminal Neuralgia. iScience. 2020;23(10):101552.

69. Baron R. Peripheral neuropathic pain: from mechanisms to symptoms. Clin J Pain. 2000;16(2 Suppl):S12–20.

70. Dhandapani R, Arokiaraj CM, Taberner FJ, Pacifico P, Raja S, Nocchi L, et al. Control of mechanical pain hypersensitivity in mice through ligand-targeted photoablation of TrkB-positive sensory neurons. Nat Commun. 2018;9(1):1640.

71. Defaye M, Iftinca MC, Gadotti VM, Basso L, Abdullah NS, Cumenal M, et al. The neuronal tyrosine kinase receptor ligand ALKAL2 mediates persistent pain. J Clin Invest. 2022;132(12).

72. Valtcheva MV, Copits BA, Davidson S, Sheahan TD, Pullen MY, McCall JG, et al. Surgical extraction of human dorsal root ganglia from organ donors and preparation of primary sensory neuron cultures. Nat Protoc. 2016;11(10):1877–88.

73. Ray PR, Shiers S, Caruso JP, Tavares-Ferreira D, Sankaranarayanan I, Uhelski ML, et al. RNA profiling of human dorsal root ganglia reveals sex differences in mechanisms promoting neuropathic pain. Brain. 2023;146(2):749–66.

74. Shiers S, Klein RM, and Price TJ. Quantitative differences in neuronal subpopulations between mouse and human dorsal root ganglia demonstrated with RNAscope in situ hybridization. Pain. 2020;161(10):2410–24.

75. Cazade M, Bidaud I, Lory P, and Chemin J. Activity-dependent regulation of T-type calcium channels by submembrane calcium ions. Elife. 2017;6.

